# A Cardiac Transcriptional Enhancer is Repurposed During Regeneration to Activate an Anti-proliferative Program

**DOI:** 10.1101/2023.06.30.547239

**Authors:** Anupama Rao, Andrew Russell, Jose Segura-Bermudez, Charles Franz, Rejenae Dockery, Anton Blatnik, Jacob Panten, Mateo Zevallos, Carson McNulty, Maciej Pietrzak, Joseph Aaron Goldman

## Abstract

Zebrafish have a high capacity to regenerate their hearts. Several studies have surveyed transcriptional enhancers to understand how gene expression is controlled during heart regeneration. We have identified *REN* or the ***r***unx1 ***en***hancer that during regeneration regulates the expression of the nearby *runx1* gene. We show that *runx1* mRNA is reduced with deletion of *REN* (Δ*REN)* and cardiomyocyte proliferation is enhanced in both Δ*REN* and Δ*runx1* mutants only during regeneration. Interestingly, in uninjured hearts, Δ*REN* mutants have reduced expression of *adamts1*, a nearby gene that encodes a Collagen protease. This results in excess Collagen within cardiac valves of uninjured hearts. The Δ*REN* Collagen phenotype is rescued even by Δ*runx1* mutations, suggesting that in uninjured hearts *REN* regulates *adamts1* independently of *runx1*. Taken together, this suggests that *REN* is rewired from *adamts1* in uninjured hearts to stimulate *runx1* transcription during regeneration. Our data point to a previously unappreciated mechanism for gene regulation during zebrafish heart regeneration. We report that an enhancer is rewired from expression in a distal cardiac domain to activate a different gene in regenerating tissue.

## Introduction

Zebrafish have a profound ability to regenerate damaged heart muscle. Heart regeneration proceeds via the proliferation of pre-existing cardiomyocytes (CMs) that replace myocardium lost to injury (Jopling et al., 2010; Kikuchi et al., 2010). Thousands of genes change expression levels in a coordinated, injury-responsive program that leads to CM proliferation and other key cell behaviors. Several groups have identified tissue regeneration enhancer elements (TREEs) in zebrafish, regulatory sequences that activate gene expression in different cell types within the regenerating heart and in other organs in response to injury and during regeneration (Cao et al., 2022; Goldman et al., 2017; Kang et al., 2016; Lee et al., 2020; Sun et al., 2022; Thompson et al., 2020). In each case, enhancer activity was demonstrated using ectopic reporter genes. However, understanding enhancer biology in its endogenous environment is more challenging because regeneration phenotypes rarely result after enhancer deletions (Sun et al., 2022; Wang et al., 2020). Phenotypes are likely difficult to recover because genes are rarely dependent on a single enhancer and multiple enhancers increase the possibility of functional redundancy (Bolt and Duboule, 2020; Kvon et al., 2021). Some examples of enhancer related regeneration phenotypes do exist and in each case these loss-of-function experiments have uncovered new and interesting biology (Sun et al., 2022; Zlatanova et al., 2023). For example, in killifish, deletion of an enhancer upstream from *inhibin beta* resulted in impaired heart regeneration and suggested a model for how evolutionary changes in enhancer elements develop together with regeneration capacity (Wang et al., 2020). However, more enhancer deletions will be necessary to reveal mechanistic activities at the core of how endogenous enhancers function during regeneration.

We produced an atlas of nucleosome turnover in zebrafish CMs to identify regeneration specific genes and regulatory elements (Goldman et al., 2017). Using a transgenic histone H3.3, we profiled changes in chromatin accessibility in CMs. Dozens of loci containing enrichment of H3.3 were validated in transgenic reporter assays as enhancers, some of which were exclusive to CMs and others that also stimulated reporter expression in other cell-types. One such reporter contained a 1265bp region found 103kb upstream from the *runx1* locus with H3.3 enrichment that increased during heart regeneration. This *runx1*-linked enhancer (or *REN*) activated GFP in proliferating cells within the wound of a regenerating heart. *REN* was not just injury responsive but was also activated in proliferating CMs in a transgenic model of CM hyperplasia where the Nrg1 protein stimulates proliferation rather than injury (Goldman et al., 2017). Thus, H3.3 profiling in regenerating hearts identified *REN* as a genetic marker for proliferating CMs in the adult. Interestingly, the *REN* enhancer is perfectly functional in mammalian hearts indicating that its regulatory machinery is likely conserved (Yan et al., 2022). However, we still have not identified what gene *REN* regulates endogenously and the importance that interaction has to regeneration.

One critical question that is incompletely understood is where regeneration enhancers come from. In part, the regeneration genetic program involves reactivation of embryonic genetic elements, for example, promoter sequences for the cardiac transcription factor *gata4* (Kikuchi et al., 2010). Afterall, regeneration is largely a recapitulation of development (Goldman and Poss, 2020; Viragova et al., 2024). However, in the case of the heart, it is unclear how much of the developmental program is necessary. The transcription factor Klf1 is dispensable for embryogenesis but required for regeneration (Ogawa et al., 2021). Moreover, Klf1 expression alone is sufficient to cause massive cardiomyocyte hyperplasia by binding mostly to enhancers that are already present in the adult (Ogawa et al., 2021). However, the enhancer(s) most critical to Klf1-driven CM hyperplasia have yet to be identified. Previously, we showed that during regeneration there is a doubling of enhancers in CMs with tens of thousands of previously inaccessible enhancers emerging. The origin of these novel CM enhancers and their importance to heart regeneration remains unknown.

Here, we demonstrate that the role of the *REN* enhancer unexpectedly changes between uninjured hearts and those that are regenerating. *REN* stimulates gene expression in a subdomain of cardiac tissue that surrounds valves leading to the outflow track of the heart. During regeneration, activity around the valve is inversely correlated with *REN* stimulation in regenerating tissue. Surprisingly, zebrafish mutants with *REN* deleted have *improved* proliferation of CMs after injury and we show that during regeneration *REN* controls the nearby *runx1* gene whose mutants have similar phenotypes. However, in uninjured hearts, *REN* deletion mutants have increased Collagen within cardiac valves. This phenotype is complemented by *runx1* mutants demonstrating that valve phenotypes are independent of *runx1*. Instead in uninjured hearts and around valves, *REN* controls *adamts1*, a metalloprotease that can degrade Collagen. Taken together, this suggests that *REN* is an enhancer that is repurposed from one cardiac domain to stimulate expression from a different gene in regenerating tissue.

### The *REN* enhancer directs gene expression in cardiac muscle and epicardium

Previously, we showed that a transgenic reporter containing the *REN* enhancer cloned upstream of a minimal promoter was able to activate GFP in proliferating cells during heart regeneration (Goldman et al., 2017). *REN* was first identified using a CM-specific profiling method which is consistent with *REN:GFP* expression co-staining with an antibody targeting the myosin heavy chain in heart muscle 7days-post-amputation (dpa) (Fig. 1A). However, we also observe a significant amount of GFP that is not in the muscle from other cardiac cell-types (Fig. 1B). The epicardium, a cell layer enveloping the heart, and endocardium, a second single-cell layer covering the heart lumen are important tissues required for signaling to the myocardium during regeneration (Kikuchi and Poss, 2012; Kikuchi et al., 2011a; Lowe et al., 2019; Lowe et al., 2021; Wang et al., 2015). To determine whether *REN:GFP* expression also occurs in the epicardium we injured fish containing both *REN:GFP* and *tcf21:dsRed* reporter transgenes and found extensive colocalization in every injured heart (Fig. 1CD) (Kikuchi et al., 2011b). Similarly, we tested *REN:GFP* and *kdrl:mCherry* double reporter fish for *REN* directed expression in endocardium (Wang et al., 2010). *REN:GFP* activates in a few isolated endocardial cells and not in every heart (Fig. 1E). Only a few isolated *REN:GFP* and *kdrl:mCherry* colocalized cells are detected within the injury area representing less than 1% of the total *kdrl*-positive cells (Fig. 1F, mean 3dpa = 2.0, mean 7dpa = 2.3; N=9,10). We conclude that *REN* is a regulatory element with predominantly myocardial and epicardial enhancer activity during heart regeneration.

**Figure 1.**
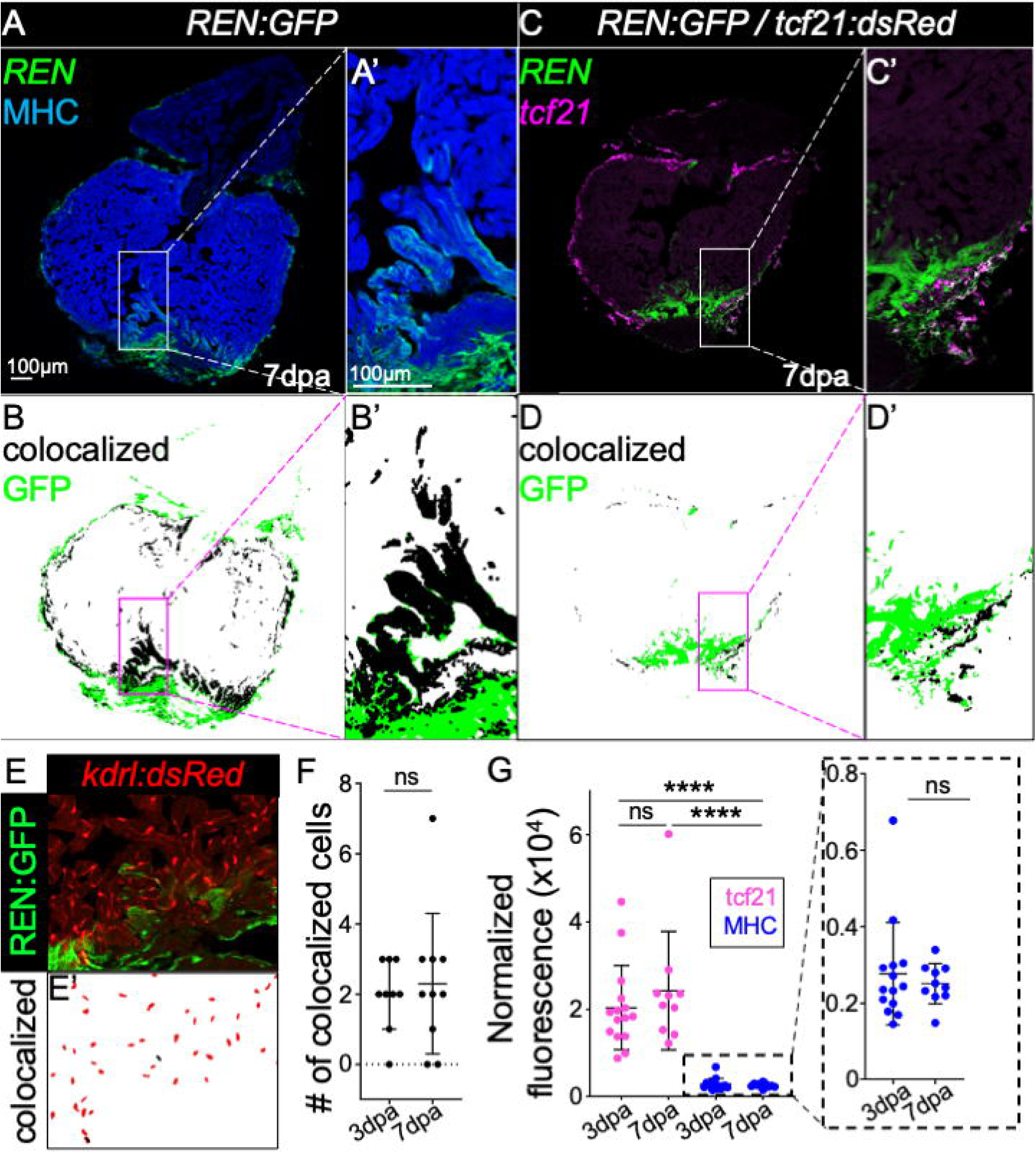
The REN enhancer expresses in epicardium and myocardium preceding the peak of regeneration. (A) Myocardial expression of *REN:GFP* is shown by staining with Anti-MHC (blue) in hearts 7-days-post-amputation (dpa). (B) MIPAR rendition of colocalization in (A) shows colocalized regions in black and GFP that is not muscle positive in green. (C) Epicardial expression of *REN:GFP* is shown by colocalization with a *tcf21*:red (pink) reporter in hearts 7dpa. (D) MIPAR rendition of colocalization in (C) shows colocalized regions in black and GFP that is not *tcf21*-positive in green. (E) Endocardial expression of *REN:GFP* is shown by colocalization with a *flk*:red reporter in hearts 7dpa. (E’) MIPAR rendition of flk colocalization from E is shown in black and *flk*-positive cells that are GFP-negative are shown in red. (F) Graph showing numbers of *flk*-positive/GFP-positive cells at 3dpa and 7dpa. (G) Graph showing the normalized GFP fluorescence of *REN:GFP* colocalized *tcf21*:Red (pink) and muscle (blue) at 3dpa and 7dpa. Zoom of muscle data is shown in the dashed box on the right.

Activation of epicardium (1-3dpa) precedes the peak of CM proliferation (7dpa) during zebrafish heart regeneration (Kikuchi and Poss, 2012). To determine the temporal regulation of *REN* driven expression we performed a time course on *REN:GFP* reporter fish. The peak of total *REN:GFP* expression occurs at 3dpa coinciding with activated epicardium throughout the heart (Fig. S1A), and is slightly decreased at 7dpa plateauing by 14-30dpa (Fig. S1B). Increased expression of GFP at 3dpa likely reflects a broader distribution of *REN:GFP* positive cells. To measure the potency of transcriptional stimulation we calculated average fluorescence intensity of GFP in colocalized cells. *REN:GFP* intensity did not change in *tcf21:dsRed* cells that colocalized with *REN:GFP* between 3 and 7dpa despite the fact that there were fewer cells at 7dpa (Fig. 1G, magenta dots, 3dpa: tcf21+GFP average = 20,395 Arbitrary Density Units(ADU)/pixel^2^; 7dpa: tcf21+GFP average = 24,269 ADU/pixel^2^; Welch’s t-test, p = 0.4481). Nor is there any change in average fluorescence intensity in CMs that colocalized with *REN:GFP* (Fig. 1G, blue dots, 3dpa:MHC+GFP average = 2770 ADU/pixel^2^, 7dpa:MHC+GFP average = 2515 ADU/pixel^2^, Mann-Whitney test, p > 0.9999). At 3dpa there is 7.36-fold more GFP intensity in epicardial cells versus CMs and 9.65-fold more GFP intensity at 7dpa, suggesting REN has more potent activity in the epicardium (Fig. 1G, 3dpa: Mann-Whitney p < 0.0001 N = 15,14, 7dpa: Mann-Whitney p < 0.0001, N=10,10). However, we cannot exclude that the observed intensity differences are impacted by cell-specific qualities such as size or shape that may influence how GFP is localized. We conclude that *REN:GFP* peaks throughout the heart at 3dpa, is focused at the site of injury by 7dpa, and is brighter in the epicardium than the myocardium.

### Minimal components of *REN* contain binding motifs for known pro-regeneration transcription factors

To identify the minimal sequence components that promote *REN* activity we produced new transgenic reporter lines with sub-fragments of *REN* cloned upstream of a minimal promoter driving GFP. Previously, we found DNA sequence motifs enriched in regeneration specific H3.3 peaks and ranked them by their specificity to regeneration (Goldman et al., 2017). Using the FIMO analysis tool we searched the *REN* enhancer for enrichment of these Cardiac Regeneration Motifs (or CRMs) and found four clusters of CRMs that included some of the most regeneration specific sequences (Fig. S2A)(Cuellar-Partida et al., 2012). Using the four clusters of CRMs as a guide, we divided *REN* into four blocks and made transgenic reporters containing these blocks in different combinations (Fig. 2A).

**Figure 2:**
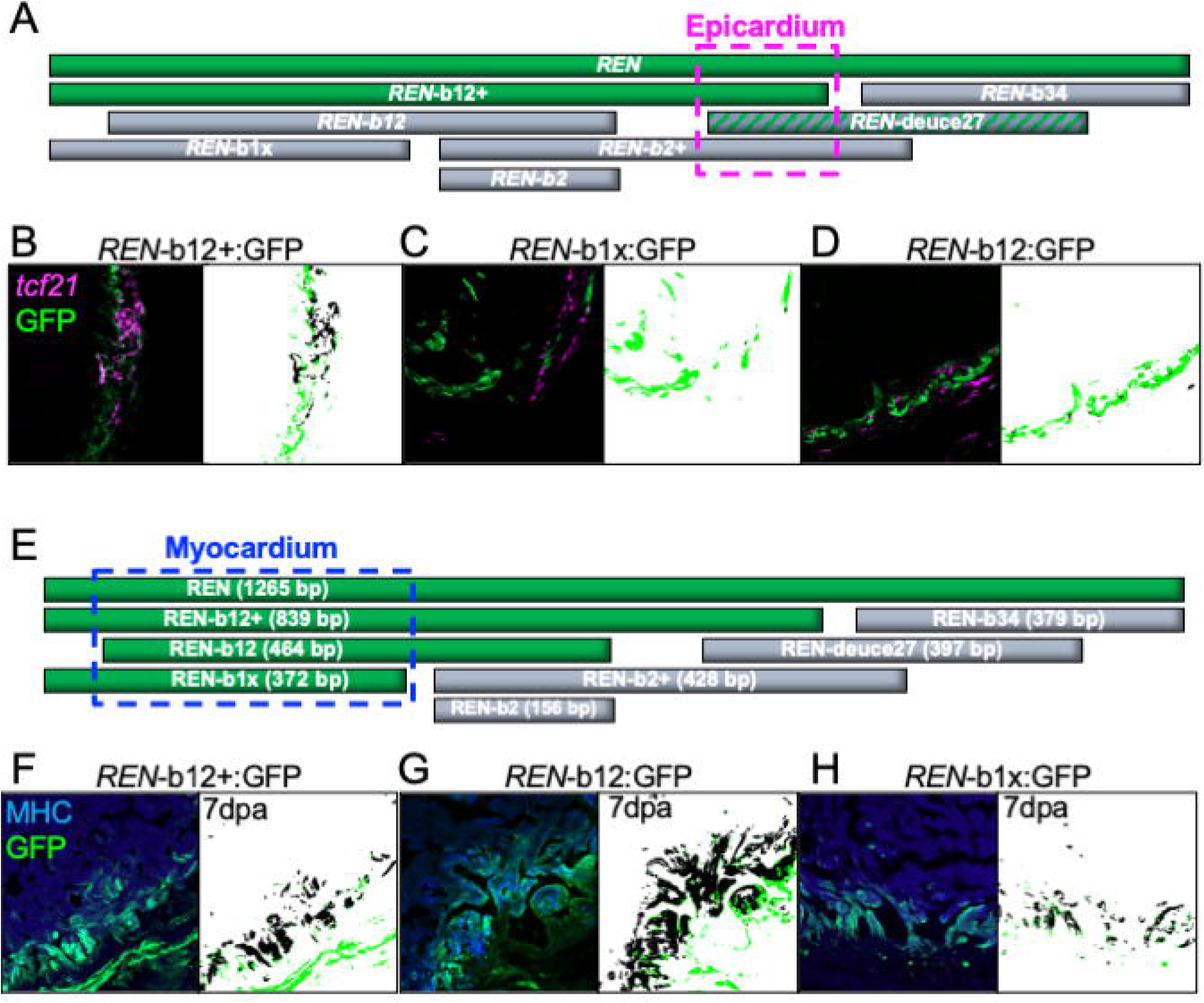
Myocardial *REN-*directed gene expression is separable from other cell types. (A) Cartoon represents a map of full-length REN (top green bar) and the seven smaller REN fragments that we tested in stable transgenic reporters. Fragments that induced GFP in epicardium are colored green and fragments that did not induce gene expression from a minimal promoter are colored gray. The epicardial specific region is outlined in a pink dashed box. (B-D) Left – Hearts from the one positive REN-b12+ transgenic and the two negative (C) REN-b12 and (D) REN-b1x reporters at 3-days-post-amputation. Right – MIPAR renditions of colocalized areas are shown in black with excess GFP remaining in green. (E) Same cartoon as in (A) except fragments of REN are colored green based on their colocalization with cardiac muscle (stained with a myosin heavy chain (MHC) specific antibody) 7dpa. (F-H) Left - Hearts from the three positive reporter lines (F) REN-b12+, (G) REN-b12, and (H) REN-b1x Right – MIPAR renditions of colocalized areas are shown in black and white with excess GFP remaining in green.

To define a minimal fragment responsible for *REN* epicardial expression we looked for fragments of *REN* that retained expression in epicardium. A fragment containing an extended region encompassing the first two blocks, called *REN*-b12+ is sufficient for activating GFP in *tcf21* positive cells (Fig. 2B). A second fragment containing just the blocks 1 and 2 called *REN*-b12 is not able to activate in the epicardium (Fig. 2C) nor is the smaller *REN*-b1x fragment (Fig. 2D). Therefore, the 375bp extended region of REN-b12+ beyond blocks 1 and 2 is necessary for epicardial activity. However, a transgenic reporter containing block 2 and the 375bp extended region (b2+) does not activate GFP at all during regeneration. This suggests that the 375bp region is necessary but not sufficient for epicardial GFP activation.

Amputation of the ventricle apex is a standard injury model that is sufficient for full-length REN activity. However, it is possible that more severe injury models would uncover GFP activity from the sub-fragments of *REN* that do not activate after amputation. Therefore, we crossed our *REN* fragments into the zebrafish cardiac ablation transgenic system (ZCAT) where CM-specific Cre releases the diphtheria toxin stochastically throughout the heart resulting in ablation of up to 60% of CMs (Wang et al., 2011). Most of the *REN* sub fragments that are silent by amputation remain silent during genetic ablation (*REN*-b34, *REN*-b2, *REN*-b2+, Fig. S2B-D). The one fragment that does activate GFP, *REN*-deuce27, does so in non-CM cells within the heart wall (Fig. S2E). The ZCAT transgenic ablation system includes a red transgenic marker so we could not confirm whether these cells are epicardial using our *tcf21* reporter. However, the expression pattern is highly reminiscent of epicardium raising the possibility that the region responsible for *REN* expression in epicardium is a 140bp fragment that overlaps with *REN*-deuce27 and *REN*-b12+. Further investigation will be required to confirm whether this 140bp region directs epicardial expression from *REN*.

The minimal fragment required for CM expression from *REN* was more straightforward. The three transgenic fragments containing block 1 can drive GFP in CMs at 7dpa (Fig. 2E-H). Any fragment of *REN* that does not have block 1 also does not activate GFP in CMs (Fig. S2B-D). The most regeneration specific motif within block 1, CRM17, has homology to binding sites of the AP1 transcription factor complex (JASPAR, p = .0175) previously reported to be required for CM proliferation (Beisaw et al., 2020). Therefore, we isolated a 267bp minimal fragment of REN that is both necessary and sufficient for *REN* myocardial activity during regeneration and that contains binding sites for known regulators of CM proliferation.

### *REN* regulates *runx1* expression in CMs and epicardium during regeneration

Multiple lines of evidence suggest that *REN* promotes a pro-regenerative gene expression program. First, *REN* directs expression in proliferating CMs independent from injury (Goldman et al., 2017). Second, *REN* activity peaks just before and during the peak of CM proliferation during regeneration (Fig. 1). Finally, a minimal fragment of *REN* that retains activity in CMs also harbors binding sites for transcription factors already shown to be required for regeneration (Fig. 2). To address whether *REN* is part of a pro-regeneration gene regulatory network, we stably deleted a 3,162bp region of the genome encompassing *REN* using CRISPR that we call Δ*REN* (Fig. S3A). We note that the 3,162 deletion encompasses the entire 1265bp of the reporter region. There is no reported evidence of other *cis*-regulatory elements being located within the 3,162bp in adult zebrafish cardiac tissues (Cao et al., 2022; Cordero et al., 2024; Goldman et al., 2017). First, we tested Δ*REN* mutants for their ability to complete regeneration. Based on the evidence from the reporter, we hypothesized that homozygous mutants lacking the *REN* enhancer would have impaired regeneration. We amputated the apex hearts from Δ*REN* mutants and their wildtype siblings and observed if regeneration was completed by 30 days. Surprisingly, the Δ*REN* mutants have complete regrowth of cardiac muscle and absence of scar that is indistinguishable from their wildtype siblings (Fig. S3BC). We conclude that *REN* is not necessary to complete regeneration.

To examine if Δ*REN* mutants have delayed regeneration, we measured CM proliferation levels at the peak of regeneration in Δ*REN* mutants and their wildtype clutch mates. We co-stained hearts recovering from amputation of the ventricle apex with Mef2, a marker for CM nuclei, and EdU, a marker of cell cycling (Fig. 3A). Unexpectedly, we find that Δ*REN* mutants have a significant *increase* in CM cycling compared to wildtype clutch mates (Fig. 3B). The fraction of Mef2 positive CMs that are also positive with the proliferation marker EdU is increased by 28.5% at 7dpa (Fig. 3B, mean: wildtype = 9.11% and mutant = 11.71%, p-value = 0.0094, N = 24 vs 19). There is no difference in CM cycling levels between Δ*REN* mutants and wildtype siblings in uninjured hearts (Fig. S3D). Also, there is no increase in overall CM numbers in adult Δ*REN* mutant hearts compared to their wildtype siblings (Fig. S3E). Thus, we conclude that Δ*REN* mutants have increased levels of CM proliferation that are specific to heart regeneration. This is in stark contrast to our expectations and suggests that *REN* is inducing expression of an anti-proliferative program.

**Figure 3:**
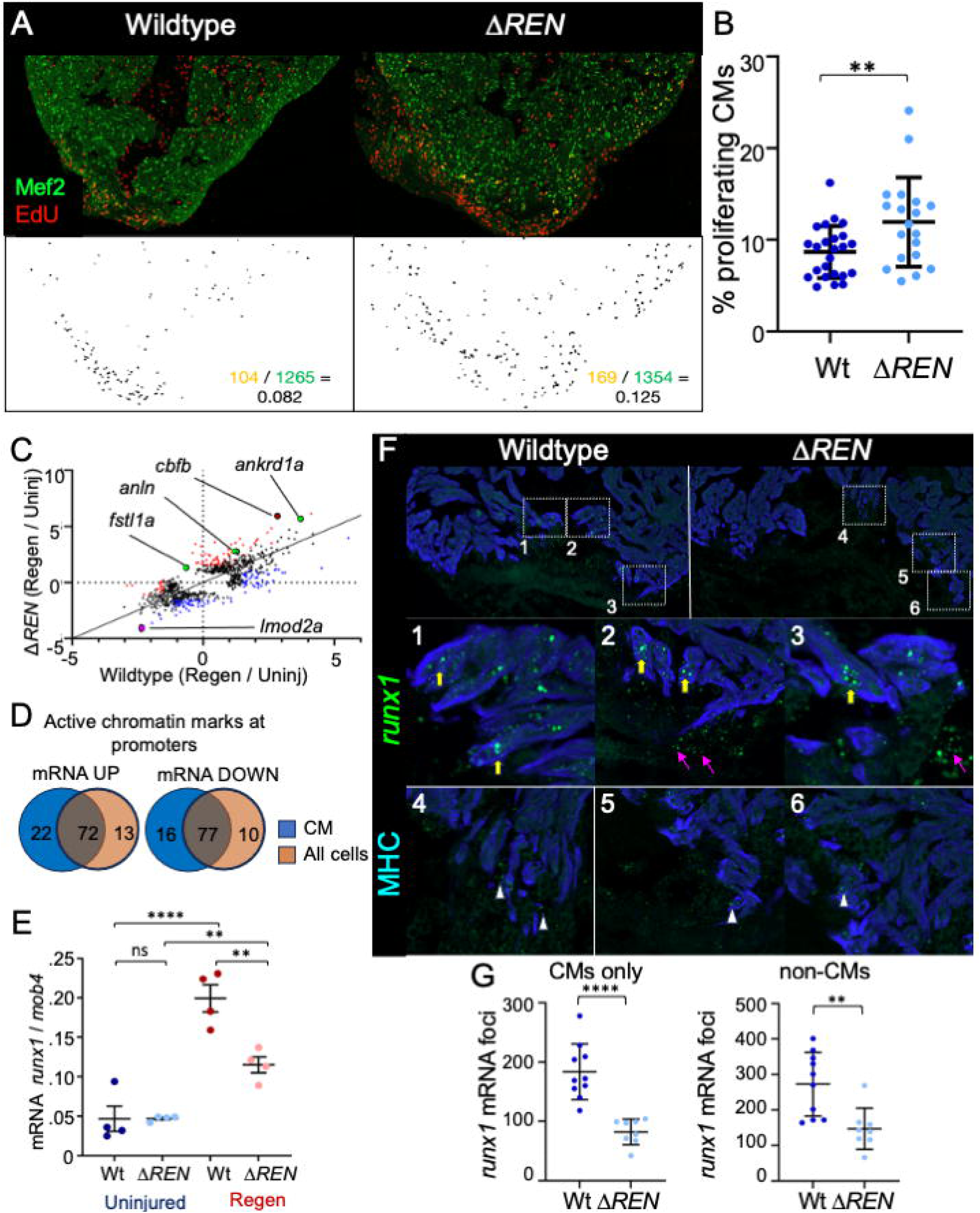
Deletion of *REN* increases cardiomyocyte proliferation during regeneration. (A) Images of sectioned amputated ventricles (7dpa) from wildtype and Δ*REN* mutant fish. Sections are stained for Mef2c (green) and EdU (red). Below – double positive cells are highlighted in black using a MIPAR software rendition (Scale bar = 100μm). (B) Quantification of CM proliferation indices (Mef2/EdU double positive over total Mef2 positive) in 7dpa ventricles (average WT = 9.11% and 11.71%; Mann-Whitney t-test, **p-value = 0.0094, N = 24 vs 19). Wildtype – blue, mutant – light blue. Horizontal black bars display the mean (middle) or standard error (top and bottom). (C) Scatterplot of RNAseq results comparing wildtype (x-axis) and Δ*REN* mutant (y-axis) samples. Each dot represents a transcript and are plotted by the log_2_ for the ratio of normalized reads from regeneration over normalized reads from the uninjured samples. Red dots are those transcripts that deviate by a linear regression >3-fold and blue dots are those transcripts that deviate by linear regression <-3-fold. Transcripts that are highlighted in the text are additionally marked in black circles. Green for pro-regeneration/proliferation genes, pink for sarcomeric genes, and runx1-coregulatory factor *cbfb*, dark red. (D) Venn diagram comparing chromatin marks at the promoters of genes whose mRNA either increases (left) or decreases (right) in Δ*REN* mutant hearts during regeneration. (E) Droplet Digital PCR shows the abundance of *runx1* transcripts increasing from uninjured wildtype hearts (blue) during heart regeneration (red) (average WT = 9.11% and MUT = 11.71%; Mann-Whitney t-test, **p-value = 0.0094, N = 24 vs 19). In Δ*REN* mutant fish (lighter colors), *runx1* levels increase less so (average WT = 9.11% and 11.71%; Mann-Whitney t-test, **p-value = 0.0094, N = 24 vs 19). Y-axis is the calculated *runx1* mRNA numbers normalized to calculated number of *mob4* mRNA. (F) Images of sectioned amputated ventricles (3dpa) from wildtype and Δ*REN* mutant fish. Sections are stained by RNAScope using a probe for *runx1* (green) and muscle was immuno-stained with an antibody towards myosin heavy chain (blue). Boxes 1-6 show a zoom of regions around the wounds highlighting runx1 mRNA within the muscle (yellow arrow), epicardial cells (pink arrow), and likely endocardial cells adjacent to muscle that remains in the mutant (white triangle). (G) The number of *runx1* mRNA foci from images like in (F) were counted using MIPAR. Quantification of foci that colocalized with muscle (MHC – Myosin heavy chain) is shown on the left (average WT = 183.1 and MUT = 82.13; Welch’s t-test, **** p-value < 0.0001, N = 10 vs 8). Quantification of foci that are not muscle is shown on the right (average WT = 272.6 and 147; Welch’s t-test, **p-value = 0.0035, N = 10 vs 8).

Since *REN* is a transcriptional enhancer, we expected that its deletion would cause gene expression changes during regeneration. Bulk RNA sequencing of Δ*REN* mutant hearts and their wildtype siblings identified transcripts that were relatively increasing or decreasing in Δ*REN* mutants during regeneration. We performed linear regression analysis on the 2,036 transcripts changing during regeneration in wildtype hearts and the 2,159 transcripts changing during regeneration in the mutant and found 305 transcripts where the fold-change during regeneration was significantly different (Fig. 3C, Table 1). There are 141 transcripts where the fold-change is relatively decreased in Δ*REN* mutants (Fig. 3C, blue dots) and another 164 transcripts where the fold-change is relatively increased in the Δ*REN* mutants (Fig. 3C, red dots). We conclude that deletion of *REN* results in dysregulation of gene expression during heart regeneration.

To identify cell-type specific expression of the differential genes we looked for enrichment of CM-specific H3.3 at their promoters (Goldman et al., 2017). Of the 305 transcripts with differential fold-change, 187 had the CM-specific H3.3 in their promoters during regeneration suggesting that the genes are expressed in CMs (Fig. 3D). The promoters of another 23 genes expressing transcripts whose levels change in Δ*REN* mutants have active chromatin marks (H3K27ac) but not H3.3-CM (Goldman et al., 2017) suggesting expression in other cell-types besides CMs. Taken together, 61% of transcripts disrupted in Δ*REN* mutant hearts are likely expressed in CMs with only 7.5% specific to other cell-types. We cannot exclude that H3.3 positive genes are being expressed in other cells *in addition* to the muscle but do conclude that upon deletion of *REN*, dysregulated genes largely occur in CMs.

Several of the 50 transcripts that increase during regeneration in Δ*REN* mutants are known regulators of cardiac regeneration. For example, in mice epicardial expression of Fstl1 increases CM proliferation and in Δ*REN* mutants *fstl1a* mRNA is 4X more abundant during regeneration (increased 22.14–fold in Δ*REN* mutants, increased 5.48-fold in wildtype)(Wei et al., 2015). Also, *fibronectin 1a* (*fn1a*) is required for zebrafish heart regeneration and *fn1a* transcripts increases 11.57–fold in Δ*REN* mutants, but only 6.12–fold in wildtype siblings (Wang et al., 2013). Expression of *ankdr1a*, also known as CARP, increases 52.82-fold in Δ*REN* mutants but only 12.96-fold in wildtype regeneration. An enhancer for CARP called *2ankrd1aEN* was found together with *REN* using H3.3 profiling and *2ankrd1aEN* can drive gene expression at the site of injury in both mouse and porcine hearts (Yan et al., 2022). Finally, there is *anln* which encodes a protein required for cytokinesis (increased 6.19–fold in Δ*REN* mutants, increased 2.41–fold in wildtype, residual = 1.72)(Oegema et al., 2000; Takayama et al., 2003). The increased induction of pro-regeneration and pro-proliferation genes supports our observation of improved proliferation of CMs (Fig. 3D) in Δ*REN* mutants.

Based on the expression of the transgenic reporter line, genes that are a direct target(s) of *REN* would be expected to *decrease* in abundance when *REN* is deleted. There were 103 transcripts that decreased relatively in mutant hearts during regeneration (Fig. 3C, red dots, Table 1). Only three of these transcripts are encoded in genes on chromosome 1 with *REN*; however, it is unlikely that they are direct targets. For example, *meis1* is transcription factor involved in maturation of CMs (Mahmoud et al., 2013), that increases 5.67-fold in wildtype regeneration but doesn’t increase in Δ*REN* mutants. However, the *meis1* gene is 49.97Mbp away from REN on the other end of the chromosome. Enhancers have been reported to interact with promoters at incredible distances including between different chromosomes (Markenscoff-Papadimitriou et al., 2014), although by chromosome capture, promoters and enhancers interact within 1Mb ∼80% of the time (Rao et al., 2014). From experimentally validated enhancer-gene associations, the largest reported distance between an enhancer and its *cis*-regulated promoter is 1.7Mb away for the Myc gene in mouse (Bahr et al., 2018; Gasperini et al., 2019). We find it unlikely that the three genes downregulated on chromosome 1 in *REN* mutants are direct targets of *REN* from 25-50 Mbp away and conclude that RNAseq alone is insufficient to identify the direct target(s) of *REN*.

Published experiments using Hi-C have detailed topologically associating domains (TADs) of self-associating chromatin from adult zebrafish brain and skeletal muscle (Yang et al., 2020). Cis-chromatin interactions within TADs helps delineate enhancer-gene pairing from chromosome looping which is remarkably similar between cell and tissue-types (Rao et al., 2014), developmental stages (Yang et al., 2020) and even between species (Harmston et al., 2017; Kikuta et al., 2007). In both zebrafish brain and skeletal muscle, the first TAD at the end of chromosome 1, encompasses *runx1* and *REN* in a 1.28Mbp domain (Yang et al., 2020) suggesting they may interact (Fig. S3F, Table 2). Activity by contact modeling of single-cell ATAC-sequencing from adult zebrafish brain (Yang et al., 2020) shows that *REN* and the *runx1* promoter are accessible within the same cells and therefore predicts *REN* to be an enhancer of *runx1* (Table 2).

To determine if *REN* regulates *runx1* in regenerating hearts, we measured changes in *runx1* mRNA in Δ*REN* mutants using the more sensitive droplet digital PCR (ddPCR) assay. For ddPCR, tens of thousands of individual PCR reactions are performed in parallel within separate lipid vesicles. The fraction of droplets that fluoresce from a successful PCR reaction are used to then extrapolate the original number of transcripts using the Poisson distribution. Using ddPCR we observed a 4.3-fold increase in the number of *runx1* transcripts during regeneration of wildtype hearts (Fig. 3E). Mutant Δ*REN* hearts, however, showed only a 2.4-fold increase or 57% of the wildtype during regeneration (Fig. 3E). The remaining increase in *runx1* abundance in Δ*REN* mutant hearts may come from endocardium, where the *runx1*-BAC reporter is induced (Koth et al., 2020) but *REN* regulates little to no expression (Fig. 1F). Therefore, we suggest that *REN* is an enhancer that regulates *runx1* expression in most cell types but not likely in the endocardium. Interestingly, mRNA for *cbfb*, a binding partner and cofactor required for Runx1 transcriptional activity, increases during regeneration in our RNAseq of Δ*REN* mutant hearts (Fig. 3C, 5.2-fold, p-value 2.84 x 10^-4^, Table 1). Likely, a feedback loop is activated without the presence of *runx1* but only during regeneration and not in uninjured hearts.

To determine the cell-type specific distribution of *runx1* expression we used RNAScope and a *runx1*-specific probe on wildtype and Δ*REN* mutant hearts (Fig. 3F). Co-staining of the *runx1* probe with an antibody towards the myosin heavy chain demonstrated that *runx1* mRNA decreases 65% in CMs of Δ*REN* mutant hearts (Fig. 3G, mean: wildtype = 183.3 and mutant = 82.13, p-value < 0.0001, N = 10 vs 8). We note that the remaining signal in the muscle may largely be background (see Material and Methods). We could not find a co-staining strategy that worked for epicardium or endocardium with RNAScope. However, non-CM signal from *runx1* mRNA also decreased 56% suggesting that *REN* controls expression in either epicardium or endocardium (Fig. 3G, mean: wildtype = 272.6 and mutant = 147, p-value = 0.0035, N = 10 vs 8). Due to the reporter expression pattern (see Fig. 1) we predict that it is in epicardial cells. The disappearing non-CM *runx1* mRNA occurs in cells invading into the wound in way similar to what is reported in the literature for epicardial cells (Fig. 3F, box 2 and 3 wildtype vs box 5 and 6 Δ*REN*)(Lepilina et al., 2006). In contrast, some of the remaining *runx1* mRNA is localized adjacent to cardiac muscle in regions reminiscent of endocardial cells (Fig. 3F, white triangles boxes 4, 5 and 6). We conclude that *REN* is a transcriptional enhancer of *runx1* expression in CMs and another cell-type(s) that is likely epicardium.

Previously, it was reported that *runx1* mutants, like Δ*REN* mutants, have increased CM proliferation during heart regeneration supporting a role for *REN* enhancing *runx1* expression (Koth et al., 2020). A major conclusion of Koth et al. was that *runx1* regulates CM proliferation through expression in endocardial cells that regulate the composition of temporary scar left within the clot after injury (Koth et al., 2020). Some of the remaining *runx1* mRNAs in Δ*REN* mutant hearts likely come from runx1 expression in endocardial cells (Fig. 3). We do not observe similar changes in the abundance of collagen and fibrin in our Δ*REN* mutants. AFOG staining of Δ*REN* mutant hearts is indistinguishable from their wildtype siblings at either 3dpa (Fig. S3G) or 7dpa (Fig. S3H). We calculated fibrin and collagen within the scars using the described methodology and while we can detect expected differences between scarring at 3dpa and 7dpa, there are no differences between Δ*REN* mutants and their wildtype siblings at any time points (Fig. S3I, 3dpa chi-square p-value = 0.378, Fig. S3J, 7dpa chi-square p-value = 0.825). These data suggest that the observed differences in scarring in *runx1* mutants are not the sole reason for observed increases in CM proliferation (please see Discussion for more details). *REN* controlled regulation by *runx1* in the epicardium and/or myocardium also affects CM proliferation. We note that the abundance of *collagen 12a1b*, which was previously reported to be pro-regenerative (Hu et al., 2022), increases only 11X during wildtype regeneration and 31X during regeneration of Δ*REN* mutant hearts (Table 1). Thus, it’s possible that *runx1* may regulate CM proliferation in epicardium or muscle in part by changing the composition of the scar.

### *REN* expression around the outflow tract of uninjured hearts is inversely correlated to expression at the site of injury

Expression of GFP in *REN:GFP* reporters is not unique to regeneration. In uninjured hearts, we observe GFP expression in CMs surrounding the valves leading from the ventricle to the outflow tract (Fig. 4A). Similar expression was observed in each of the three independent *REN* reporter lines that we originally generated (Goldman et al., 2017). As seen during regeneration, *REN:GFP* also expresses in non-CMs including in epicardial cells that emanate above the valve around the outflow tract (Fig. 4B). Calculation of relative GFP intensities showed that *REN:GFP* has 10X more GFP around uninjured valves as the rest of the uninjured ventricle (Fig. S4A) Interestingly, *REN:GFP* expression disappears in the myocardium surrounding the valve during regeneration with its nadir at 7dpa, it begins to return by 14dpa (Fig. 4C-F). At 30dpa, when most of the muscle has finished regrowing (Fig. S3A), *REN:GFP* expression reaches an equilibrium where it is on slightly less than half of its maximal expression in CM both around valves and in the freshly regrown site of injury (Fig. 4G). The expression of *REN* recovers to an uninjured distribution by 60dpa (Fig. 4G, yellow line), however at lower overall levels (Fig. 4G, red and blue lines). In summary, the peak of *REN* regulated gene expression is inversely correlated between CMs adjacent to valves and CMs that are undergoing regeneration. This suggests that *REN* activity is mechanistically connected between the expression domains.

**Figure 4:**
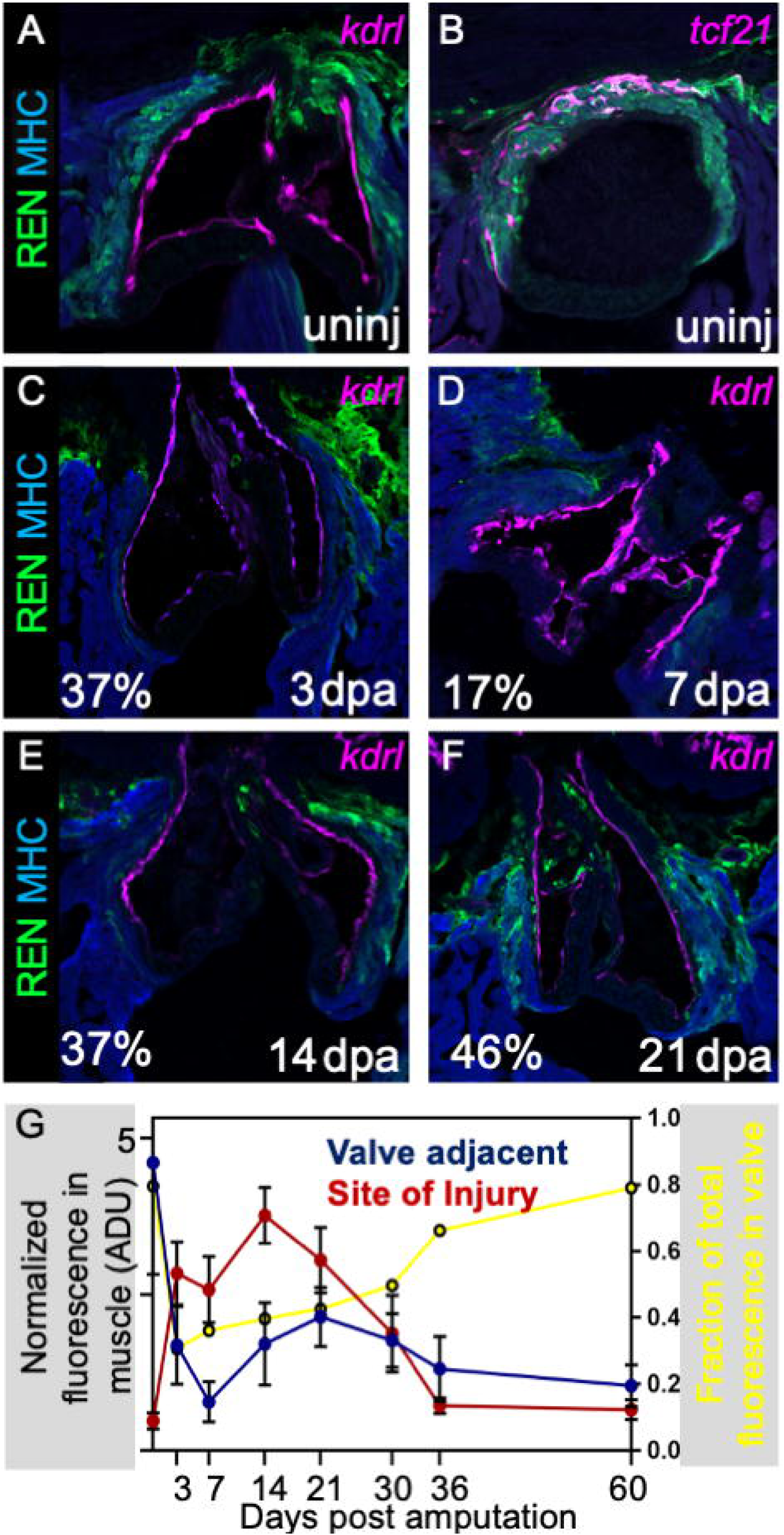
*REN* regulates gene expression in cardiac tissue surrounding valves in uninjured hearts. (A) Valves around the outflow tract in uninjured hearts are shown by endocardial reporter *flk*:Red (pink). *REN:GFP* (green) and muscle is stained with MHC (blue). (B) both cardiac muscle (MHC-blue) and epicardium (*tcf21* reporter, pink) colocalize with *REN:GFP* (green) around uninjured valves. (C-F) Same staining as in (A) but this time on regenerating hearts 3dpa (C), 7dpa (D), 14dpa (E), and 21dpa (F). Percentage of remaining GFP intensity is shown in the bottom left corner. (G) Quantification of average EGFP intensity in regions colocalizing with cardiac muscle (MHC). Cardiac valve regions are shown in blue, and site of injury is shown in red. X-axis is a timeline for the days after amputation of the ventricle apex. Y-axis (left) is the normalized fluorescence. Y-axis (right) is the fraction of the total fluorescence that is found around valves (yellow line).

To determine the minimal component of *REN* required for expression around valves, we looked at *REN* fragment reporter expression in uninjured hearts. *REN* fragments b12+ (Fig. S4A) and b1x (Fig. S4B) both have activity in myocardium surrounding valves in uninjured hearts and the b2+ fragment does not (Fig. S4C). Thus, the 267bp minimal region of *REN* that is necessary and sufficient during regeneration is also necessary and sufficient for myocardial expression in uninjured hearts. We conclude that the same region of the genome is repurposed to activate genes in cardiac tissue, between uninjured valves and during regeneration.

### Deletion of *REN* causes gene expression changes affecting outflow tract valves

On occasion, reporters for transcriptional enhancers have subdomains of ectopic expression that do not reflect endogenous expression patterns (Bolt and Duboule, 2020). To confirm that the endogenous *REN* locus regulates genes in cardiac tissue adjacent to valves, we analyzed our RNAseq comparing Δ*REN* mutants and their wildtype siblings only looking at the uninjured replicates. There are 158 transcripts whose abundance significantly decreases by 75% and another 137 transcripts whose abundance significantly increases at least 3-fold in uninjured hearts of Δ*REN* mutants (Fig. S5A, Table 3). The differential abundance of mRNA transcripts suggests that *REN* does indeed regulate gene expression *in vivo* in uninjured hearts. Likely, many of the transcripts changing in abundance are not direct targets and are secondary effects of *REN* disruption. Interestingly, gene ontology analysis of transcripts with increased abundance in Δ*REN* mutants included three members of the AP1 transcription factor complex, *fosab*, *fosb* and *atf3* each of which increase 26.3-fold, 11.62-fold and 9.08-fold respectively in uninjured mutant hearts (Fig. S5A, green dots). Therefore, disruption of *REN* resulted in increased levels of mRNAs encoding a transcription factor complex with motifs present in the deleted enhancer (Fig. S2A). This suggests that a feedback loop may be in place regulating *REN* activity in uninjured hearts.

The *REN:GFP* reporter shows that *REN* activates genes, suggesting that a direct target of *REN in vivo* would be among the transcripts that are less abundant in uninjured Δ*REN* mutant hearts. Of the 158 less abundant transcripts, 6 are encoded by genes found on chromosome 1 and none are found within the reported TAD with *REN* where most enhancer-promoter interactions lie (Sun et al., 2019; Symmons et al., 2014) (Fig. S3F, Table 2). However, another gene called *adamts1,* 672Mb away from *REN* did have decreased abundance (Table 1). The *adamts1* gene encodes an extracellular matrix protein that regulates cardiac valves by degrading collagen (Hong-Brown et al., 2015). We used RNAscope to determine changes in the spatial distribution of *adamts1* mRNA in wildtype and Δ*REN* mutant hearts. As expected, expression of *adamts1* mRNA was strong within endocardial interstitial cells that compromise cardiac valves (Fig. 5A, asterisks). It was also abundant in neighboring CMs (Fig. 5A, blue arrow) and likely epicardial cells (Fig. 5A, yellow arrow). In Δ*REN* mutant hearts, the abundance of *adamts1* transcripts decreased 62% in CM surrounding valves (Fig. 5B, mean: wildtype = 81.1and mutant = 31.1, Mann-Whitney p-value = 0.0001, N = 8 vs 9). Expression of *adamts1* also decreased to 66% in non-CM cells (Fig. 5C, mean: wildtype = 955.4 and mutant = 321.8, Mann-Whitney p-value = 0.0015, N = 9 vs 10). Therefore, we conclude that *REN* functions as a transcriptional enhancer for the *adamts1* gene in CMs and possibly in the epicardium surrounding cardiac valves. The expression of *adamts1* also increases at the site of injury during regeneration (Table 1). Interestingly, there was no significant difference between *adamts1* abundance in CMs at the site of injury between wildtype and Δ*REN* mutant hearts (Fig. 5C, mean: wildtype = 28.0 and mutant = 23.9, Welch’s p-value = 0.634). The average level of *adamts1* mRNA is decreased 40% in epicardial cells although the p-value is not significant (Fig. 5C, mean: wildtype = 48.6 and mutant = 29.5, Welch’s p-value = 0.088). We conclude that *REN* is not a critical enhancer for inducing *adamts1* in regenerating tissue but it’s possible that *REN* fplays a more minor role as a shadow enhancer that stabilizes expression of *adamts1* in nonCMs.

**Figure 5:**
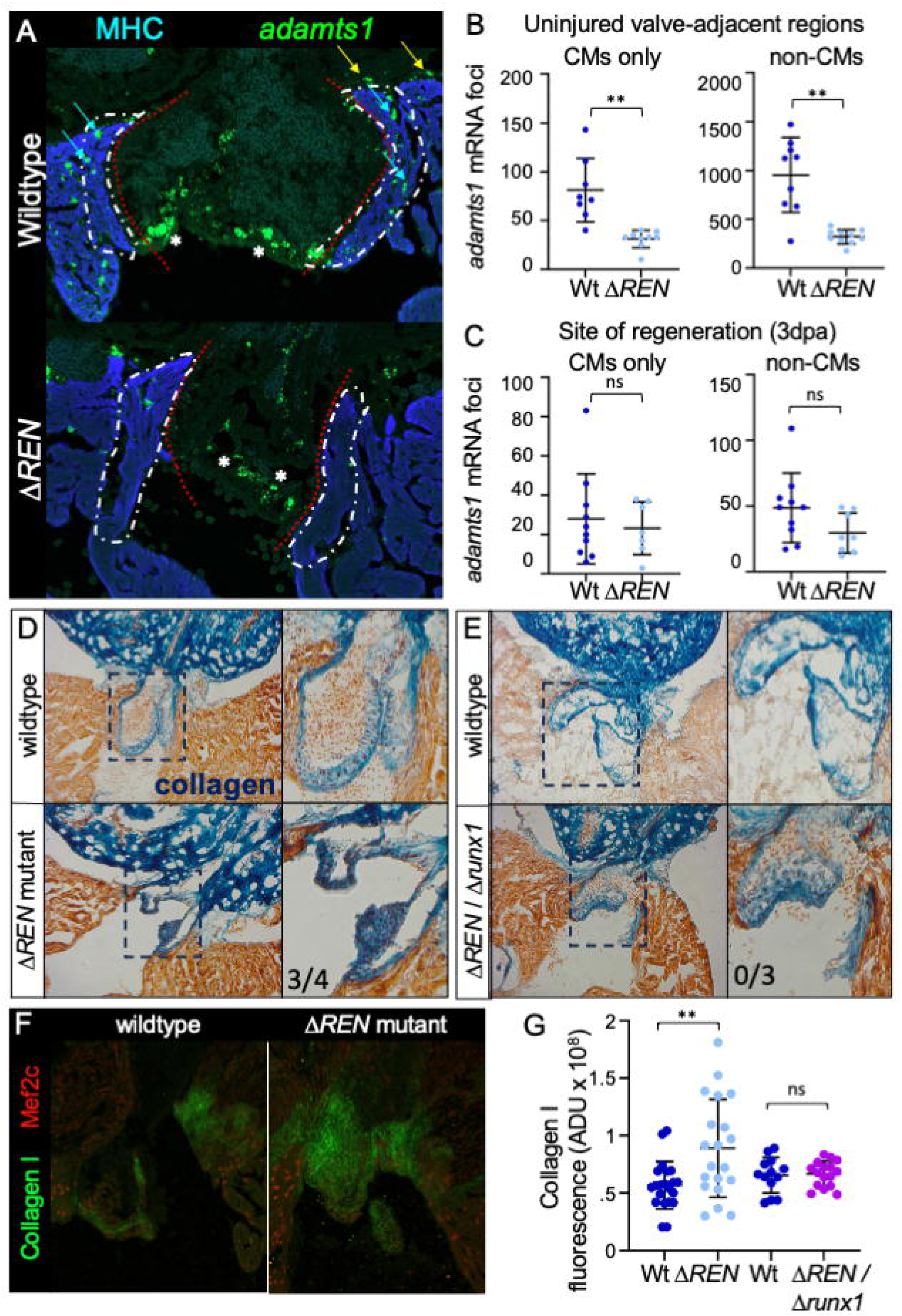
*REN* regulates different genes in uninjured hearts and during regeneration. (A) Images of uninjured valves from wildtype and Δ*REN* mutant fish. Sections are stained by RNAScope using a probe for *adamts1* (green) and muscle was immuno-stained with an antibody towards myosin heavy chain (blue). (B) The number of *adamts1* mRNA foci from images like in (A) were counted using MIPAR. Quantification of foci that colocalized with muscle (MHC – Myosin heavy chain) is shown on the left (mean: wildtype = 81.1 and mutant = 31.1, Mann-Whitney p-value = 0.0001, N = 8 vs 9). Quantification of foci that are not muscle is shown on the right (mean: wildtype = 955.4 and mutant = 321.8, Mann-Whitney p-value = 0.0015, N = 9 vs 10). Wildtype – blue, mutant – light blue. Horizontal black bars display the mean (middle) or standard error (top and bottom). (C) The number of *adamts1* mRNA foci from the sectioned amputated ventricles (3dpa) analyzed for runx1 in Fig. 3F. Quantification of foci that colocalized with muscle (MHC – Myosin heavy chain) is shown on the left (mean: wildtype = 28.0 and mutant = 23.9, Welch’s p-value = 0.634). Quantification of foci that are not muscle is shown on the right (mean: wildtype = 48.6 and mutant = 29.5, Welch’s p-value = 0.088). (D-E) AFOG staining of uninjured wildtype and uninjured Δ*REN* mutant hearts (D) and uninjured wildtype and uninjured Δ*REN* / Δ*runx1* double heterozygote hearts (E). Valve leaflets are shown in a zoom on the right. (F) Valves around the outflow tract in uninjured wildtype and Δ*REN* mutant hearts are immuno-stained with anti-Collagen I (green) and anti-Mef2c (red) to mark CM nuclei. (G) Quantification of Collagen I stain intensity (mean arbitrary density units (ADU): wildtype (blue) = 5691576603 and mutant (light blue) = 8904512333, Mann-Whitney p-value = 0.0091, N = 21 vs 20; wildtype = 6545145907 and double het (purple) = 6671972330, Welch’s p-value = 0.799, N = 13 vs 16).

To see if Δ*REN* mutants had observable phenotypes associated with Adamts1, we stained Δ*REN* mutant hearts for collagen using AFOG and compared them to their wildtype siblings (Fig. 5D, S5C). Uninjured valves surrounding the outflow tract are visibly bluer in Δ*REN* mutant hearts indicating that they have more Collagen. Several pieces of data suggest that *REN* expression in uninjured valves is distinct from *runx1* (Koth et al., 2020). First, from the *runx1*-BAC reporter, there is no detectable fluorescence in uninjured hearts (Koth et al., 2020). Second, there is no detectable difference in *runx1* expression between uninjured wildtype and uninjured Δ*REN* mutant hearts (Fig. 3F). To determine if *REN* and *runx1* are in different pathways in uninjured hearts, we performed a genetic complementation experiment (Fig. S5D). We crossed Δ*REN* heterozygotes to a *runx1* mutant line and compared compound heterozygotes containing one of each of the *REN* and *runx1* mutant alleles to their wildtype siblings. Uninjured hearts were stained for collagen by AFOG, and compound heterozygotes displayed similar levels of collagen as their wildtype siblings demonstrating that uninjured phenotypes in Δ*REN* mutant hearts do not depend on *runx1* (Fig. 5E). To quantitatively assess Collagen differences in Δ*REN* mutant valves we used immunofluorescence with an antibody towards Collagen I (Fig. 5F). In Δ*REN* mutant hearts, Collagen I increased 56% in valves (Fig. 5G, mean arbitrary density units: wildtype (blue) = 5691576603 and mutant (light blue) = 8904512333, Mann-Whitney p-value = 0.0091, N = 21 vs 20). This difference was rescued by crossing Δ*REN* to a Δ*runx1* mutant demonstrating that Collagen changes are independent of *runx1* (Fig. 5G, mean arbitrary density units: wildtype = 6545145907 and double het (purple) = 6671972330, Welch’s p-value = 0.799, N = 13 vs 16). Taken together, deletion of *REN* results in both decreased abundance of a cardiac valve associated gene, *adamts1,* and in phenotypes associated with reduced Adamts1 function. We conclude that *REN* regulates *adamts1* expression around uninjured cardiac valves.

## Discussion

Here, we describe a transcriptional enhancer called *REN* that is active in cardiac tissue around the valve in uninjured hearts but is repurposed to activate at the site of injury during heart regeneration (Fig. 4). Targeted removal of this enhancer using CRISPR resulted in increased CM proliferation during regeneration and in increased collagen levels in valve leaflets in uninjured hearts (Fig. 3 and 5). Changes in underlying gene expression together with genetic complementation experiments show that different genes are regulated by *REN* during regeneration and in uninjured hearts (Fig. 5). The activity of transcriptional enhancers during regeneration has been well characterized (Sun et al., 2022; Wang et al., 2020; Yan et al., 2022; Zlatanova et al., 2023). Yet, the source of regenerative enhancers, how they are recruited, and then subsequently targeted to specific genes is less well understood. Previously, it has been shown that regenerative enhancers reemerge from developmental enhancers or by invigoration of pre-existing adult enhancers (Kikuchi et al., 2010; Ogawa et al., 2021). In addition, here, we show that during regeneration, a particular enhancer is repurposed from functioning around cardiac valves and recruited to activate a different gene that is important for CM proliferation in regenerating tissue (Fig. 6A).

**Figure 6:**
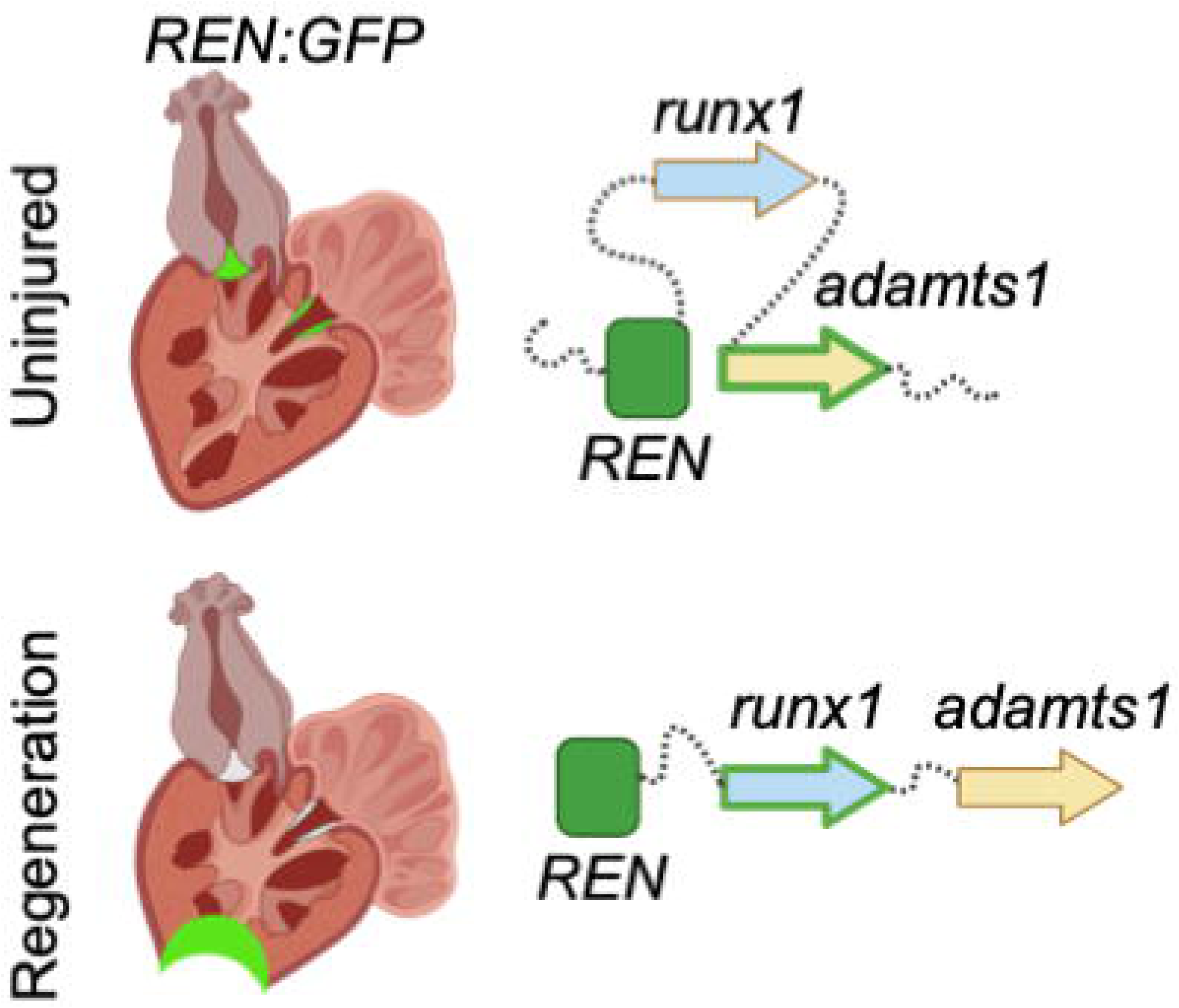
*REN* is rewired from a different cardiac domain *and* to a different gene during heart regeneration.

Several lines of evidence demonstrate that *REN* very likely directly regulates *runx1* during regeneration. First, deletion of either gene or enhancer results in a similar phenotype; CM proliferation improves during regeneration (Koth et al., 2020). Second, single cell ATAC-seq from zebrafish brains uncovered an ‘enhancer hub’ where the promoters for *runx1* and genes for calcium channel homologs *atp1a1a.4*, *atp1a1a.3*, and *atp1a1a.5* all associate with *REN* (Fig. S3F)(James et al., 1999; Uyehara and Apostolou, 2023; Yang et al., 2020). This suggests that at least in the brain, *REN* forms a ‘hub’ to co-regulate these four genes across ∼350Mbp. Third, when *REN* is deleted, *runx1* expression is decreased during regeneration (Figs. 3E and F), confirming that also in the heart, *REN* regulates *runx1* expression. There must be additional enhancer(s) that govern *runx1* expression in endocardial cells during regeneration (Fig.1F) and *runx1* expression during hematopoietic development since Δ*REN* and *runx1*^Δ*TSS*^ double heterozygous mutants develop at Mendelian ratios unlike a *runx1* mutant homozygote (Koth et al., 2020; Lam et al., 2009; Sood et al., 2010). *REN* regulates *runx1* expression in CMs during regeneration (Fig. 3G). However, from the RNAScope alone, we cannot conclusively say whether additional *runx1* mRNA changes are occurring in epicardial or in endocardial cells (Fig. 3G). The localization of the changing *runx1* mRNA suggests deletion of *REN* affects the epicardium (Fig. 3F) which is consistent with both *REN:GFP* reporter expression (Fig. 1F) and the reported data from the Koth et al. manuscript. We conclude that *REN* is a CM and epicardial specific enhancer of *runx1* during regeneration (Fig. S6A).

Around cardiac valves in uninjured hearts, *REN* has additional roles independent of *runx1.* First, unlike *REN*, *runx1* reporters do not express around uninjured cardiac valves (Koth et al., 2020). Second, *runx1* mRNA levels do not change when *REN* is deleted from uninjured hearts (Fig. 3F). Third, the *runx1* co-regulator *cbfb* is only increased during regeneration in mutants and not in uninjured hearts indicating that a feedback loop that invigorates missing *runx1* function is specific to regeneration. Finally, and most importantly, the *runx1* mutant allele compensates for Δ*REN* mutant phenotypes around valves demonstrating that valvular phenotypes are independent of *runx1* (Figs. 5E and 5G). We note that the 3,162bp deletion in the Δ*REN* mutant may remove a previously undetected *cis*-regulatory element that regulates valvular genes. However, the same minimal fragment of *REN* is necessary and sufficient to activate during regeneration is also necessary and sufficient to activate in uninjured valves (Fig. 4). Thus, the Δ*REN* deletion mutant and reporter experiments are consistent with one another suggesting the same enhancer element is driving expression in both contexts.

What is the gene that *REN* regulates in uninjured hearts if it is not *runx1*? It is possible that valvular collagen changes result from other gene(s) within in the *REN* enhancer ‘hub’ (Fig. S3F), for example, misexpression of sodium potassium ion channels like *atp1a1a* (James et al., 1999). Yet, we do not detect expression of mRNA from the cluster of *atp1a1a* homologs and there are no active chromatin marks at the three *atp1a1a* promoters in zebrafish hearts (Goldman et al., 2017). We postulate that *REN* regulates the *adamts1* gene either directly or indirectly for several reasons. First, in uninjured Δ*REN* mutant hearts, valves have more collagen, a phenotype consistent with decreased Adamts1 function (Santamaria and Groot, 2020) (Fig. 5BC). Second, while *adamts1* does not sit in the same TAD as *REN* in brain and muscle, there are interactions across the *REN* containing TAD boundary between both the *adamts1* and *runx1* promoters (Yang et al., 2020) (Fig. S6B, Table 3). The *runx1* promoter interacts with an enhancer upstream of *adamts1* and the *adamts1* promoter interacts with an enhancer within the *REN*-containing TAD (Yang et al., 2020). Third, the abundance of *adamts1* mRNA decreases in uninjured Δ*REN* mutant hearts around cardiac valves in the same region the *REN:GFP* reporter expresses (Fig. 5A). This raises the possibility that *REN* regulates *adamts1* across 672Mbp in cardiac tissue around valves in uninjured hearts.

Regeneration phenotypes for enhancers mutants are difficult to recover. There is a double requirement to have both a gene that when mutated would result in a phenotype *and* whose expression is dominated by a single enhancer. Here, we show that differences between enhancer mutants and gene mutants, can reveal new mechanistic insights of gene function. In mutants for *runx1*, Koth et al. reported changes in clot composition in the wound during regeneration that is not observed in our Δ*REN* mutants (Fig. S3G-J). Since, unlike *runx1*, *REN* does not robustly activate in endocardial cells (Fig. 1F), we surmise that observed changes in scar deposition result from perturbing endocardial *runx1* expression (Koth et al., 2020). However, the shared increases in CM proliferation in the *runx1* and Δ*REN* mutants demonstrate that *runx1* impacts CM proliferation through either epicardium or muscle. We cannot exclude a possible role for endocardial *runx1* in regulating CM proliferation as well. However, the Δ*REN* mutant clearly establishes epicardial and/or muscle expression of *runx1* as being critical for negatively regulating CM proliferation during regeneration. More work is needed to dissect the individual cell-type specific contributions and to identify direct targets of Runx1 that modify the ability to regenerate hearts.

Loss of function mutations that result in improved heart regeneration are not commonly described (Koth et al., 2020; Missinato et al., 2018). Why an anti-proliferative gene would be activated during regeneration is not obvious and poses a bit of a paradox. For example, at the peak of regeneration the *REN* enhancer is both a genetic marker of CM proliferation *and* activates an anti-proliferative program (Fig. 3) (Goldman et al., 2017). *A priori* one might expect that a program to slow regeneration would be temporally activated towards the conclusion of regeneration rather than the peak. We see no phenotypes in our Δ*REN* mutant hearts that would suggest regeneration is halted abnormally (data not shown); however, we cannot exclude that there are redundancies regulating the end of regeneration. In mice, Runx1 is induced in certain models of dilated cardiomyopathy and knockout of *Runx1* in heart muscle improves recovery after heart injury (Kubin et al., 2011; McCarroll et al., 2018). However, changes in CM proliferation levels have not been described. Recently, *Runx1* has been reported to be linked to the abundance of more proliferative mono-nucleus diploid CMs (MNDCMs), which raises the possibility that expression of Runx1 in early stages of mouse postnatal development increase numbers of MNDCMs by preventing binucleation through inhibiting cytokinesis (González-Rosa et al., 2018; Patterson et al., 2017; Swift et al., 2023).

## Acknowledgments

We would like to thank the American Heart Association for their support (GRANT #17SDG33660922 to JAG) and the Department of Biological Chemistry and Pharmacology, College of Medicine, Ohio State University Medical Center for funding. The Δ*REN* mutant strain and some *REN* fragment reporter lines were originally derived in Ken Poss’s lab, and we are grateful for his support and comments on this manuscript. Also, thanks to Arthur Burghes for access to the Droplet Digital PCR machine, loaning us Anton Blatnik, and for manuscript comments. Thanks to Maria Mihaylova and Kubra Akkaya for training with the RNAScope. Thanks to Dr. Joshua Waxman for advice on the Collagen antibody. Thank you to the Neuroscience Imaging Core at the Ohio State University (#P30-NS104177) for use of the confocal microscope.

## Material and Methods

### Transgenic fish construction

For reporter fish strains, the regulatory element being studied was cloned upstream of the mouse fos minimal promoter as previously described (Goldman et al., 2017). Stable transgenic zebrafish lines were produced using the I-Sce method of random genomic integration. The *tcf21*:Red reporter and the *flk*:Red reporters were previously published. All zebrafish (*Danio rerio*) used in this study derive from the Ekwill strain. Adults less than 1 year old were used for all the experiments. Males and females were mixed in similar proportions in each of the conditions. All experiments were performed under university supervision according to the institutional animal care and use committee protocol #2018R00000090-R2 of Ohio State University. For enhancer reporter lines, we selected at least 2 and as many as 6 independent insertions to exclude potential insertional effects.

### Heart injuries

Zebrafish were anesthetized using Tricaine and placed ventral side up on a sponge to carry out resection of the ventricular apex. Iridectomy scissors were used to make an incision through the skin and pericardial sac. Gentle abdominal pressure exposed the heart and ∼20% of the apex was removed with scissors, penetrating the chamber lumen (Poss et al., 2002). Hearts were harvested 1, 3, 7, 14, 21 or 30 days after injury depending on the experiment. To genetically ablate CMs, *cmlc2:CreER^pd10^; bactin2:loxp-mCherry-STOP-loxp-DTA^pd36^* (Z-CAT) fish were incubated in 0.5 μM tamoxifen for 17 h (Wang et al., 2011).

### Immunofluorescence

Primary antibodies used in this study were: Rabbit anti-Mef2 (1:100, Abcam, ab197070), Rabbit anti-GFP (1:200, Life Technologies A11122), Mouse anti-MF20 (1:100, Developmental Studies Hybridoma Bank, MF20), and Mouse anti-Collagen I (1:10, Developmental Studies Hybridoma Bank, SP1.D8-s). For anti-Mef2, hearts were embedded fresh frozen. For collagen, slides were boiled 5 minutes in citrate buffer for epitope unmasking using a steamer. Secondary antibodies were: Goat anti-mouse Alexa Fluor 546 (1:200, Thermo Fisher Scientific, A-11030), and Goat anti-rabbit Alexa Fluor 488 (1:200, Thermo Fisher Scientific, A-11034).

### Quantification of fluorescence

#### Average fluorescence

Briefly, using ImageJ, the red (or blue) channel was thresholded and used to select a region of interest in the green channel where the raw integrated density was calculated. Background was subtracted. The calculated fluorescence was then normalized to area under consideration.

#### CM proliferation

Injured fish were injected into the abdominal cavity once every 24 h for 3 days (4-6 dpa) with 10 μl of a 10 mM solution of EdU diluted in PBS. Hearts were removed on day 7, embedded and cryosectioned. Slides were stained with Alexa Fluor 594 Azide using click chemistry (Breinbauer and Köhn, 2003) and then immunostained for Mef2c. Briefly, sections were blocked with 1% bovine serum albumin (Fraction V) and 5% goat serum and washed in PBS with 0.2% Triton X-100. Three sections representing the largest wound area were selected from each heart and imaged using a 20× objective. The number of Mef2^+^ and Mef2^+^EdU^+^ cells were counted using MIPAR image analysis software and the CM proliferation index was calculated as the number of Mef2^+^EdU^+^ cells/total Mef2^+^ cells (Sosa et al., 2014). The CM proliferation index was averaged across two to four appropriate sections from each heart.

#### Mutant fish construction

To derive the Δ*REN* allele (os^76^), guide RNAs (sgRNAs) were designed against two regions flanking the REN element on chromosome 1 (sgUpstream: gTA gTg TTg Agg ATA gAC Ag, sgDownstream: gAA AAC AgC TAC AgC TCC CT). DNA templates of the respective sgRNA fused to a tracRNA were produced by PCR. T7-transcribed sgRNA were then injected with Cas9 protein into newly fertilized embryos from EK parents. Mutant strains were genotyped using PCR oligos (Fwd: gCC ACT gCC TCg CCC CTg Cg, Rev: AAT CgA TgA TTC TTg Agg TCA AAg ATg TgT ACT) for the mutant allele. A third oligo (ACA gCT ACA gCT CC TCg gAC) was added to the PCR to identify the wildtype allele. *REN* deletion was perfect without addition of extra nucleotides between the fused cut sites. The 3162bp deletion encompasses the entire 1265bp fragment from the reporter plus some adjacent sequence that consists of mostly transposons. Finding reliable sgRNA and genotyping primer pairs within highly repetitive regions was non-trivial.

#### RNAseq analysis

Demultiplexed and quality-filtered reads, were aligned to the Danio rerio reference genome GRCz10 using Hierarchical Indexing for Spliced Alignment of Transcripts 2 (HISAT2) (Kim et al., 2015). Read counts for each gene were quantified using featureCounts software (Liao et al., 2014). Differential gene expression analysis was performed using R package edgeR (McCarthy et al., 2012). The read counts were normalized using TMM method (Robinson and Oshlack, 2010). Differentially expressed genes were selected based on adjusted P value and log2 fold change.

#### Linear regression analysis

A master list of transcripts was created using the following criteria: (1) The transcript of interest had a P-value<0.05 for both the WT regeneration and Mut regeneration. (2) The transcript of interest had a log_2_FC>1 for either WT regeneration or Mut regeneration. Each transcript was assigned its WT regeneration log_2_FC as the X-value, and its Mut regeneration log_2_FC as the Y-value. A line of best fit was calculated using the master list, and residuals were calculated for each transcripts using the line of best fit. If its residual>1, then the transcript was colored red. If its residual<-1, then the transcript was colored blue.

#### AFOG staining

Acid Fushin Orange G staining was performed as previously described (Rao et al., 2023). For calculation of relative fibrin and collagen levels we adopted the methodology described in (Koth et al., 2020). Briefly, ImageJ was used to analyze the color of the wound area of sections stained with AFOG. Channels were split and images were color thresholded for red and blue using the same settings for all hearts. Red and blue particles were analyzed and %Area was recorded for each heart. To determine the orange area, %Red and %Blue values were subtracted from 100%. Percents were averaged from at least 5 hearts from each condition.

#### Statistics

For each analysis, we used the Welch’s parametric *t*-test to calculate significance. If the variance between the comparison groups was also significant (*F* test for unequal variances), we switched to using a non-parametric test (Mann–Whitney). For the CM counting and CM proliferation assays, one researcher embedded hearts and sectioned slides and a separate researcher who was unaware of the sample identity carried out imaging and quantification.

#### Droplet-digital PCR

mRNA was isolated from wild-type and Δ*REN* uninjured and injured hearts using Trizol (Goldman et al., 2017). cDNA was made using 1µg RNA in Superscript II reverse transcriptase reactions incubated at 44 °C for 50 minutes. ddPCR assays were developed to detect *runx1*, *adamts1*, and *mob4* cDNA. cDNA for Mob4 served as loading control as it does not change in RNA-seq datasets in response to injury or between cell-types (REF Rao and from Rao). The primer and probe sequences for ddPCR are: *runx1* forward primer 5’-CgA gAg CCA CgA CgC CAC-3’, *runx1* reverse primer 5’-CgA CTg CTC ATA CgA CCA ggA Tgg-3’, runx1 probe 5’-/56-FAM/CAT gCg gTg/ZEN/ CAg CCC ACA CCA Cg/3IABkFQ/-3’; mob4 forward primer 5’-AgT ATT TTC CCA gCC gCg TCA gC -3’, mob4 reverse primer 5’-TCA CgA AAC ggg TgA AgC gAT gAC -3’, mob4 probe 5’-/HEX/TCC CAT gCg/ZEN/ TAC TTT CAC CAT CgC CAg/3IABkFQ/-3’; adamts1 forward primer 5’-gAg ACC TgC CCT gAT AGC AAT gg-3’, adamts1 reverse primer 5’-ggT gTC CCA TCA gCC ACC T-3’, adamts1 probe 5’-/56-FAM/CTG CAA gTT/ZEN/ ggT gTG CCg AgC gAA GG/3IABkFQ/-3’. We used 1 µL of reverse transcriptase reactions for detection of *runx1* and *adamts1* cDNA, and 1 µL of a 1:10 dilution of the cDNA was used to quantify *mob4*. Standard BioRad reaction conditions were used—10 µL 2x ddPCR Supermix for Probes (No dUTP) (cat# 1863023) 0.9 µM forward primer, 0.9 µM reverse primer, 0.25 µM probe, desired template amount and remaining volume of water to achieve 20 uL reaction volume. Reactions were partitioned into droplets using BioRad QX200 Droplet Generator (cat# 1864002), mixing 20 µL of reaction with 70 µL Droplet Generation Oil for Probes (cat#1863005). Droplets were transfered to Eppendorf 96-well twin.tec semi-skirted 96-well PCR Plates (cat# 951020389) and sealed with BioRad Pierceable Foil Heat Seal (cat# 1814040) using a Vitl Life Science Solutions Variable Temperature Sealer (cat# V902001). PCR thermocycling conditions were performed as follows: 50 °C for 2 minutes, 95°C for 2 minutes, 55 cycles of 95 °C 30 seconds, 60 °C for 1 minute, and 72 °C for 30 seconds, followed by 72 °C for 30 seconds, 12 °C forever, using BioRad T100 Thermal Cycler (cat# 1861096). Reactions were read using BioRad QX200 Droplet Reader (cat# 1864003) and thresholds were drawn between the positive and negative droplet populations using the BioRad QX Software Version 2.1. Data was exported to Apple Numbers for further processing. Analysis of resulting data had to conform to >10,000 accepted reaction droplets, >1000 negative reaction droplets, and the number of positive droplets had to be greater than samples that did not receive reverse transcriptase. The calculated copies/µL for *runx1* and *adamts1* were normalized to *mob4*. These ratios were directly imported into GraphPad Prism to generate the figures.

#### RNA Scope

slides with hearts fixed in paraformaldehyde (4% o/n at 4C) were sectioned, placed at -20C for 2hrs and then moved to -80C overnight. From there we strictly followed the ACD (Biotechne) Technical Note Sample preparation for fixed from tissue using RNAscope 2.5 Chromogenic assay. For the RNAscope protocol itself we used the RNAscope Multiplex Fluorescent Reagent Kit v2 with the following modifications. We used protease IV, diluted fluorophores 1:3000, washed 3X each for 5 minutes with gentle mixing after. This was to get rid of as much background signal as possible in the muscle. We boiled slides for 5 minutes for the *adamts1* probe (made for this project) or 10 minutes for the *runx1* probe (Koth et al., 2020). After the last developing and wash for the RNAscope, we washed slides 3X in PBS with .05% Tween, left them to block 1 hr at RT in 1% goat serum, 1% BSA, and then in primary with the MHC (MF20) antibody 1:100 o/n at 4C. Slides were washed and coverslipped the next day as normal.

## Supplemental Figure Legends

**Supp Figure 1 –**

(A) Time course of *REN:GFP* expression throughout the heart during regeneration. REN – green, MHC – blue. (B) Quantification of total area containing GFP fluorescence.

**Supp Figure 2 –**

(A) Cartoon of REN with example Cardiomyocyte Regeneration Motifs (CRM) shown from each of the four blocks. (B-E) Heart sections from the different REN fragments (labeled) in ZCAT hearts ablated 7 days-post-induction. Left – gray scale of GFP. Right – MHC (red), REN:GFP (green).

**Supp Figure 3 –**

(A) Diagram of the Δ*REN* deletion mutant showing site of chromosome break and the 9 nucleotides inserted during the repair. (B) Wildtype and mutant hearts were stained with MHC 30 days-after-amputation. (C) AFOG staining of the same hearts. (D) Quantification of CM proliferation indices (Mef2/EdU double positive over total Mef2 positive) in uninjured ventricles (wildtype average = 0.78%; mutant average = 0.54%; Welch’s t-test, p-value = .252, N = 7 vs 11). Wildtype – blue, mutant – light blue. Horizontal black bars display the mean (middle) or standard error (top and bottom). (E) Graph of total Mef2-positive CM numbers counts in adult uninjured hearts (average = 2095 and 1666; Welch’s t-test, p-value = 0.115, N = 7 vs 11). (F) Cartoon of the TAD containing both *REN* and *runx1* in adult zebrafish brain and muscle. (F’) Cartoon showing the ‘*REN* enhancer hub’ where promoters for *runx1* and the three *atp1a1a* genes all interact with one another and with *REN*. Created in BioRender.com (G-H) Representative image of injury site from wildtype and D*REN* mutant hearts stained with AFOG at 3 days-post-amputation (G) and 7dpa (H). (I) Calculation of relative fibrin (red) and collagen (blue) levels from 3dpa based on Koth et al. methodology. wildtype averages: collagen = 7.75%; fibrin = 8.70%; muscle (orange) = 83.73%; mutant averages: collagen = 5.16%; fibrin = 12.60%; muscle (orange) = 82.25%; Chi-square p-value = 0.3376; N=7,5. (J) Calculation of relative fibrin (red) and collagen (blue) levels from 7dpa based on Koth et al. methodology. wildtype averages: collagen = 1.13%; fibrin = 5.36%; muscle (orange) = 93.51%; mutant averages: collagen = 1.35%; fibrin = 3.89%; muscle (orange) = 93.51%; Chi-square p-value = 0.825; N=5,6.

**Supp Figure 4 –**

(A) Minimal fragment of REN is sufficient for CM expression around uninjured valves (REN-b1X); muscle (blue) and GFP (green). Right - MIPAR rendition of colocalized areas from REN-b1X are shown in black with excess GFP remaining in green. (B) REN fragments b12+ (C) and b2+.

**Supp Figure 5 –**

(A) Volcano plot showing differences in RNAseq from uninjured wildtype hearts vs uninjured Δ*REN* hearts. Transcripts of genes decreasing within 1.7Mb of REN on chromosome 1 are highlighted in pink and labeled with arrows. Members of the AP1 transcription factor complex are highlighted in green and labeled with arrows. (B) AFOG staining of uninjured wildtype and uninjured Δ*REN* mutant hearts. Images are zoomed in on the region around valves near the outflow tract. All replicates are included here to show support Figure 5DE. (C) Cartoon of complementation experiment. Shown are the regions of chromosome 1 deleted in the Δ*REN* and Δ*runx1* mutant lines (dashed blue boxes). The Y-axis is the enrichment of cardiomyocyte specific–histone H3.3 and the X-axis are coordinates along chromosome 1. (D) Immunofluorescence of cardiac vlaves with Collagen I (green) and Mef2c (red) antibodies. replicates are included here to support Figure 5F.

**Supp Figure 6 –**

(A) The *REN* enhancer deletion effectively acts as a CM and epicardial knockout of *runx1* during regeneration. After injury *REN:GFP* only expresses in CMs and epicardium but *runx1* is also endocardial. Thus, knockout of *REN* has *wild type* endocardial expression of *runx1*. Proliferation phenotypes in ΔREN therefore arise from CMs or epicardium. (B) Cartoon of the entire region of chromosome 1 including the telomeric end and the TAD containing both *REN* and *runx1*. Genes are shown as arrows with dotted lines representing 3D interactions found by Hi-ChIP. The loop marked 1 is the interaction of the *adamts1* promoter with an enhancer within the TAD. The loop marked 2 is the interaction between the *runx1* and an enhancer outside of the TAD nearby *adamts1*. Created in BioRender.com

