## Supplemental Figures for "A Cardiac Transcriptional Enhancer is Repurposed During Regeneration to Activate an Anti-proliferative Program"

**Supp Figure 1 –**

(A) Time course of *REN:GFP* expression throughout the heart during regeneration. REN – green, MHC – blue. (B) Quantification of total area containing GFP fluorescence.

Supp Figure 1

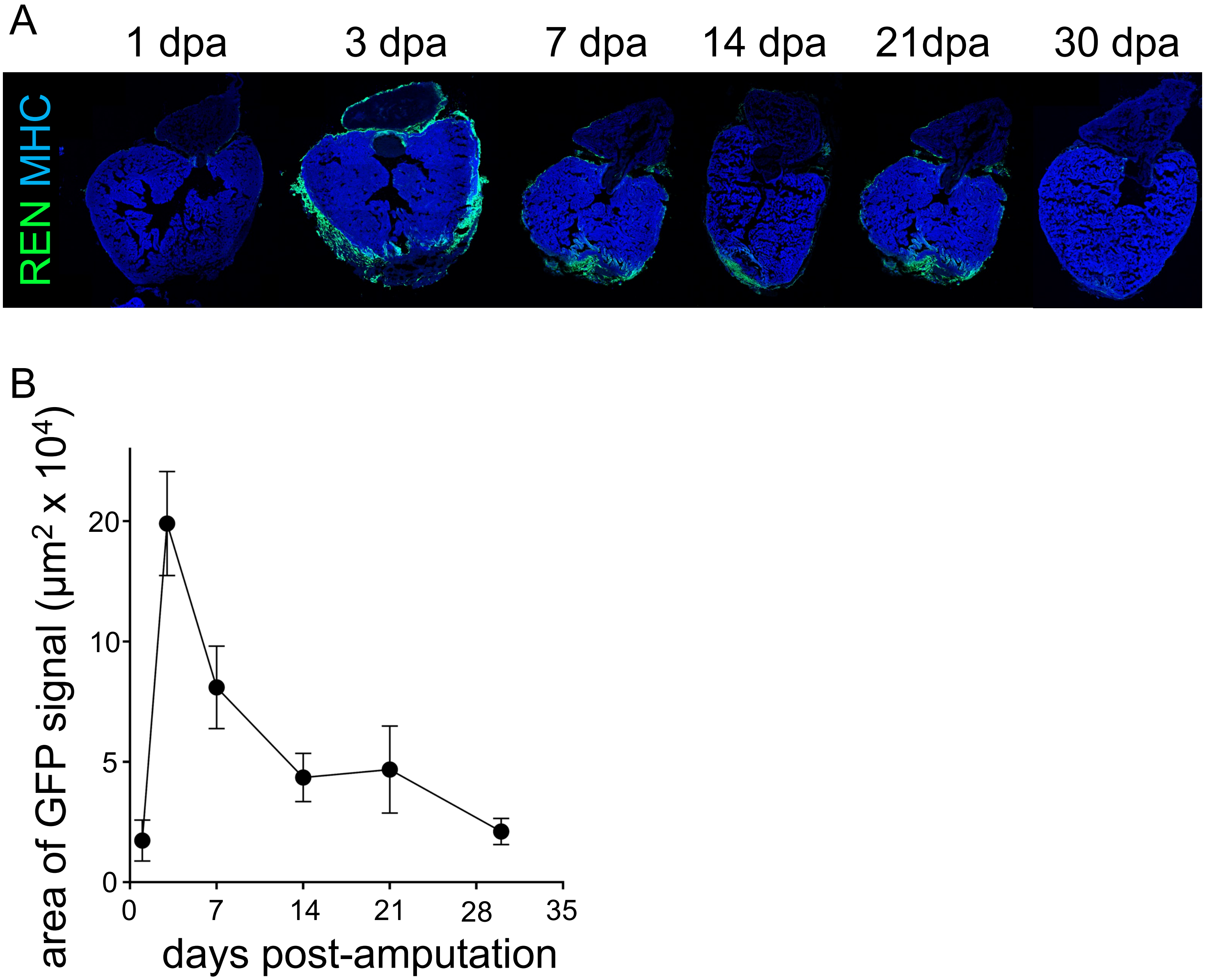

**Supp Figure 2 –**

(A) Cartoon of REN with example Cardiomyocyte Regeneration Motifs (CRM) shown from each of the four blocks. (B-E) Heart sections from the different REN fragments (labeled) in ZCAT hearts ablated 7 days-post-induction. Left – gray scale of GFP. Right – MHC (red), REN:GFP (green).

Supp Figure 2

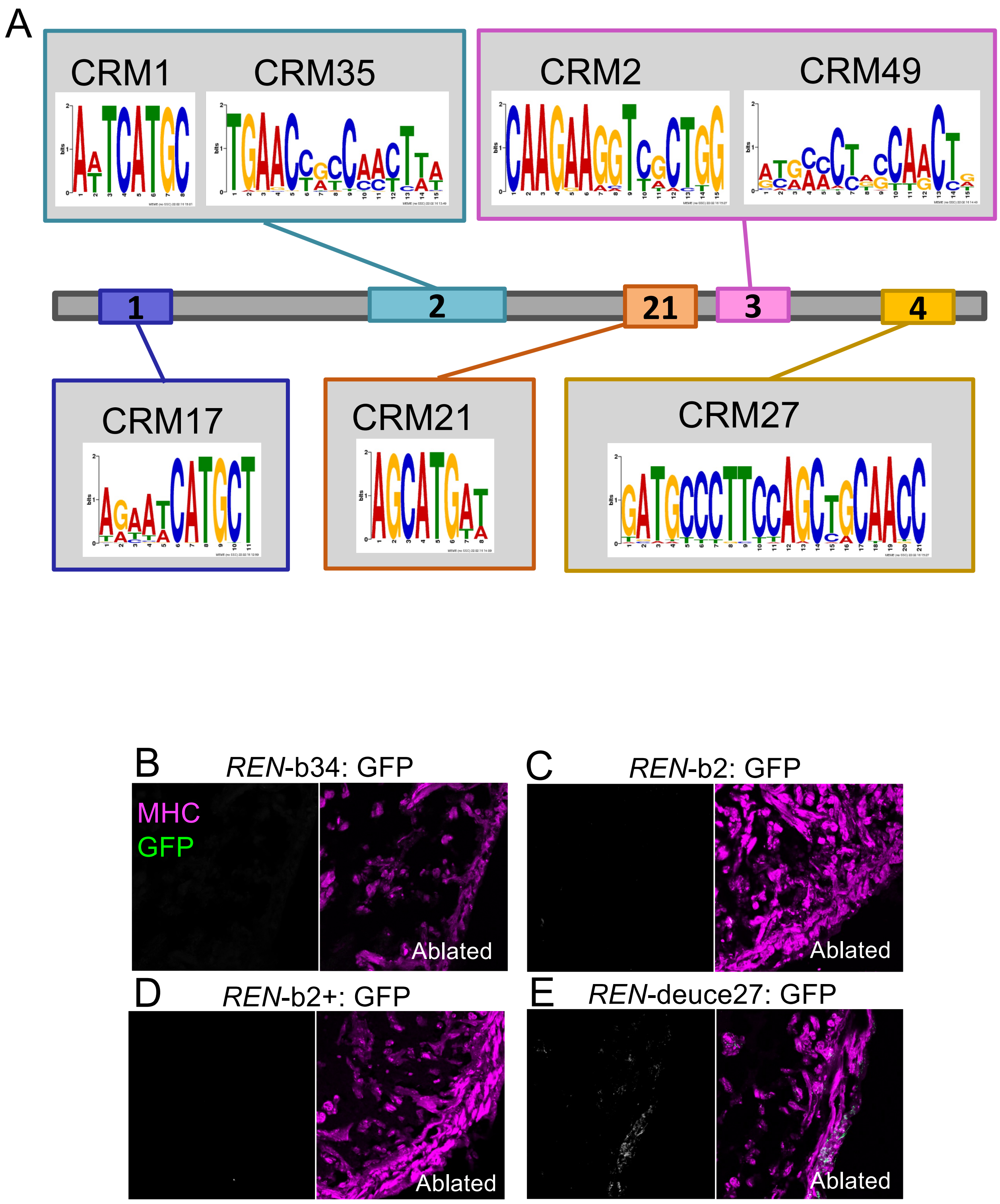

**Supp Figure 3 -**

(A) Diagram of the Δ*REN* deletion mutant showing site of chromosome break and the 9 nucleotides inserted during the repair. (B) Wildtype and mutant hearts were stained with MHC 30 days-after-amputation. (C) AFOG staining of the same hearts. (D) Quantification of CM proliferation indices (Mef2/EdU double positive over total Mef2 positive) in uninjured ventricles (wildtype average = 0.78%; mutant average = 0.54%; Welch's t-test, p-value = .252, N = 7 vs 11). Wildtype – blue, mutant – light blue. Horizontal black bars display the mean (middle) or standard error (top and bottom). (E) Graph of total Mef2-positive CM numbers counts in adult uninjured hearts (average = 2095 and 1666; Welch's t-test, p-value = 0.115, N = 7 vs 11). (F) Cartoon of the TAD containing both *REN* and *runx1* in adult zebrafish brain and muscle. (F') Cartoon showing the '*REN* enhancer hub' where promoters for *runx1* and the three *atp1a1a* genes all interact with one another and with *REN*. Created in BioRender.com (G-H) Representative image of injury site from wildtype and Δ*REN* mutant hearts stained with AFOG at 3 days-post-amputation (G) and 7dpa (H). (I) Calculation of relative fibrin (red) and collagen (blue) levels from 3dpa based on Koth et al. methodology. wildtype averages: collagen = 7.75%; fibrin = 8.70%; muscle (orange) = 83.73%; mutant averages: collagen = 5.16%; fibrin = 12.60%; muscle (orange) = 82.25%; Chi-square p-value = 0.3376; N=7,5. (J) Calculation of relative fibrin (red) and collagen (blue) levels from 7dpa based on Koth et al. methodology. wildtype averages: collagen = 1.13%; fibrin = 5.36%; muscle (orange) = 93.51%; mutant averages: collagen = 1.35%; fibrin = 3.89%; muscle (orange) = 93.51%; Chi-square p-value = 0.825; N=5,6.

Supp Figure 3

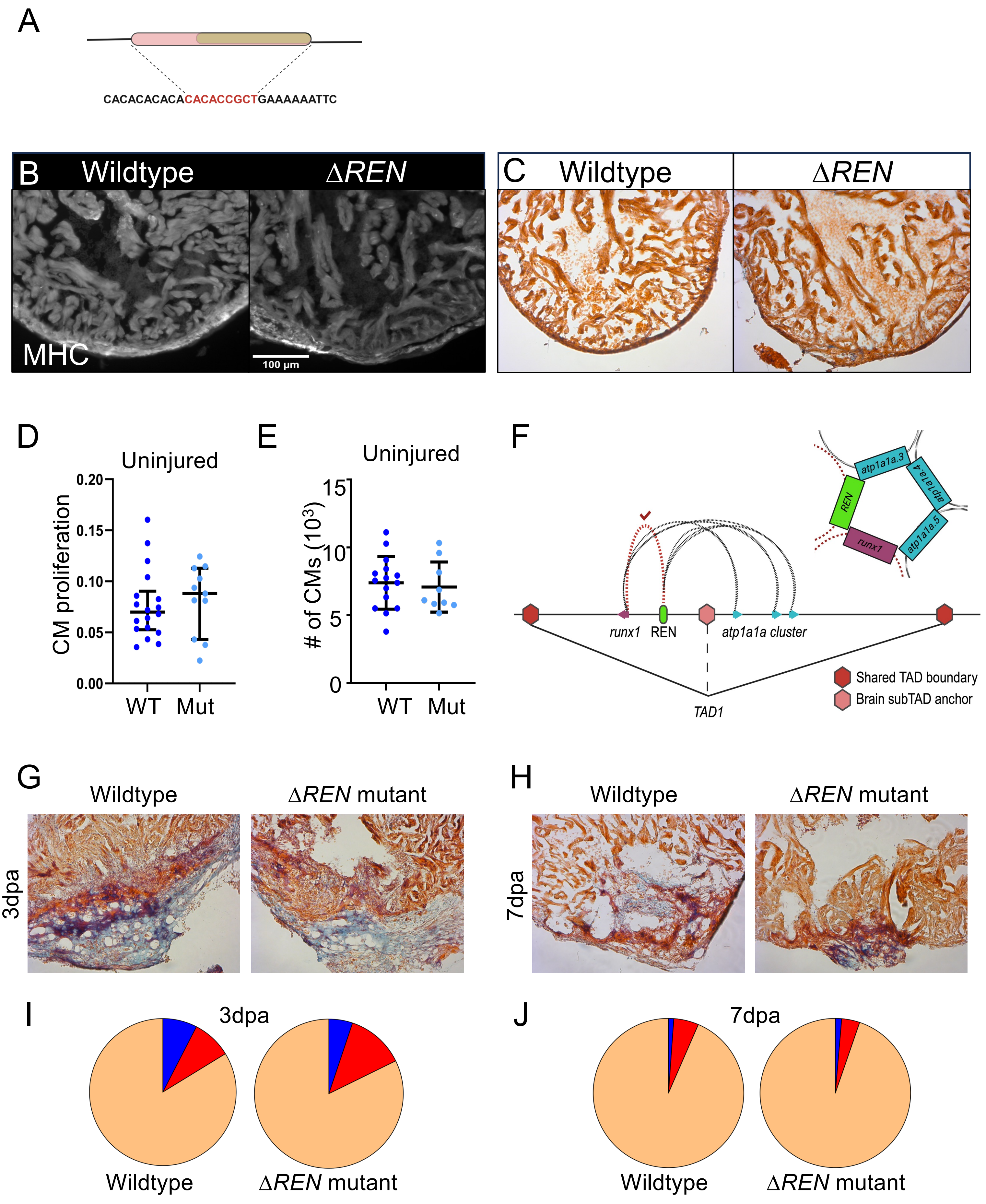

**Supp Figure 4** –

(A) Minimal fragment of REN is sufficient for CM expression around uninjured valves (REN-b1X); muscle (blue) and GFP (green). Right - MIPAR rendition of colocalized areas from REN-b1X are shown in black with excess GFP remaining in green. (B) REN fragments b12+ (C) and b2+.

Supp Figure 4

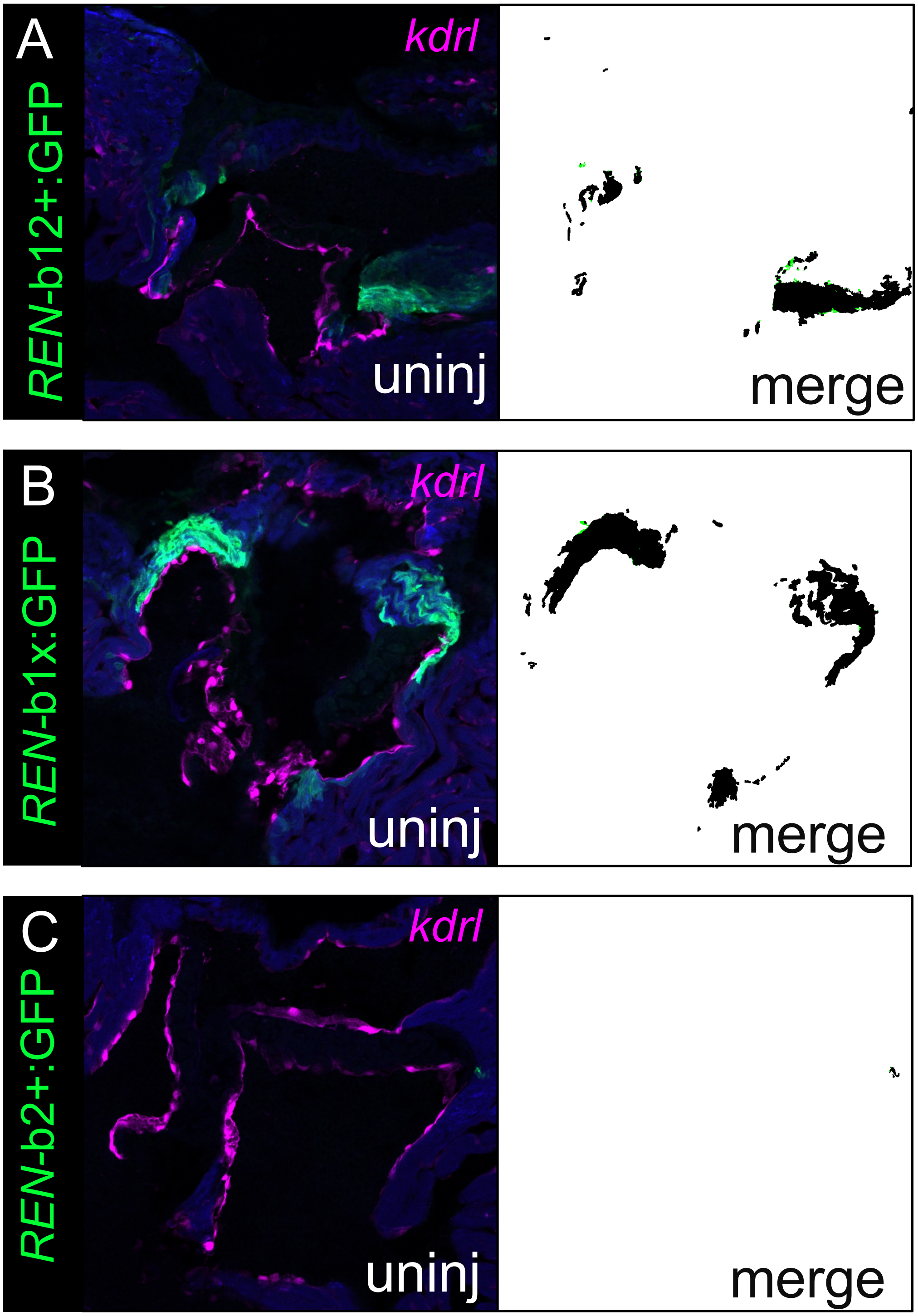

**Supp Figure 5 –**

(A) Volcano plot showing differences in RNAseq from uninjured wildtype hearts vs uninjured ∆*REN* hearts. Transcripts of genes decreasing within 1.7Mb of REN on chromosome 1 are highlighted in pink and labeled with arrows. Members of the AP1 transcription factor complex are highlighted in green and labeled with arrows. (B) AFOG staining of uninjured wildtype and uninjured ∆*REN* mutant hearts. Images are zoomed in on the region around valves near the outflow tract. All replicates are included here to show support Figure 5DE. (C) Cartoon of complementation experiment. Shown are the regions of chromosome 1 deleted in the ∆*REN* and ∆*runx1* mutant lines (dashed blue boxes). The Y-axis is the enrichment of cardiomyocyte specific–histone H3.3 and the X-axis are coordinates along chromosome 1. (D) Immunofluorescence of cardiac vlaves with Collagen I (green) and Mef2c (red) antibodies. replicates are included here to support Figure 5F.

Supp Figure 5

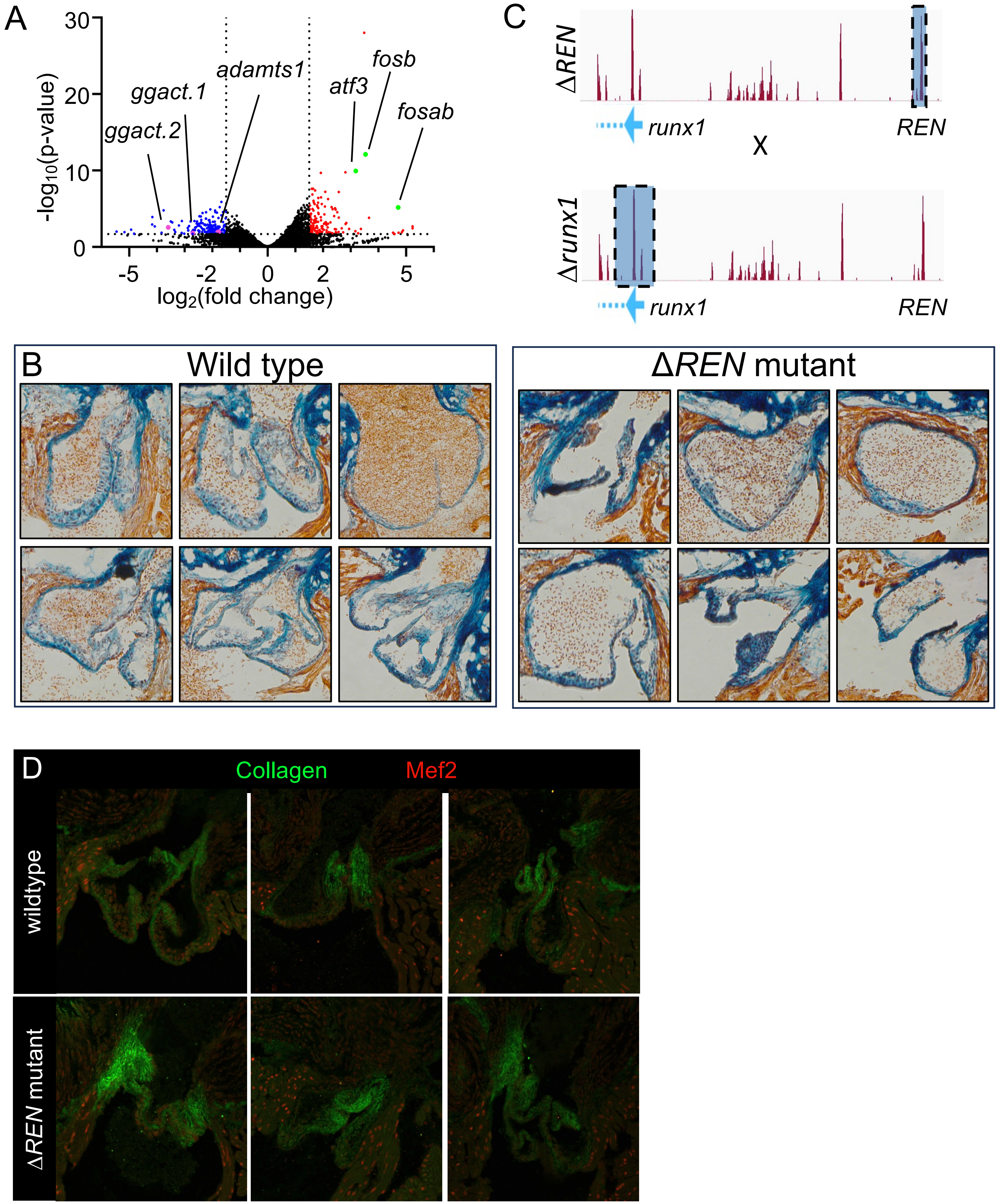

(A) The *REN* enhancer deletion effectively acts as a CM and epicardial knockout of *runx1* during regeneration. After injury *REN:GFP* only expresses in CMs and epicardium but *runx1* is also endocardial. Thus, knockout of *REN* has *wild type* endocardial expression of *runx1*. Proliferation phenotypes in ∆REN therefore arise from CMs or epicardium. (B) Cartoon of the entire region of chromosome 1 including the telomeric end and the TAD containing both *REN* and *runx1*. Genes are shown as arrows with dotted lines representing 3D interactions found by Hi-ChIP. The loop marked 1 is the interaction of the *adamts1* promoter with an enhancer within the TAD. The loop marked 2 is the interaction between the *runx1* and an enhancer outside of the TAD nearby *adamts1*. Created in BioRender.com

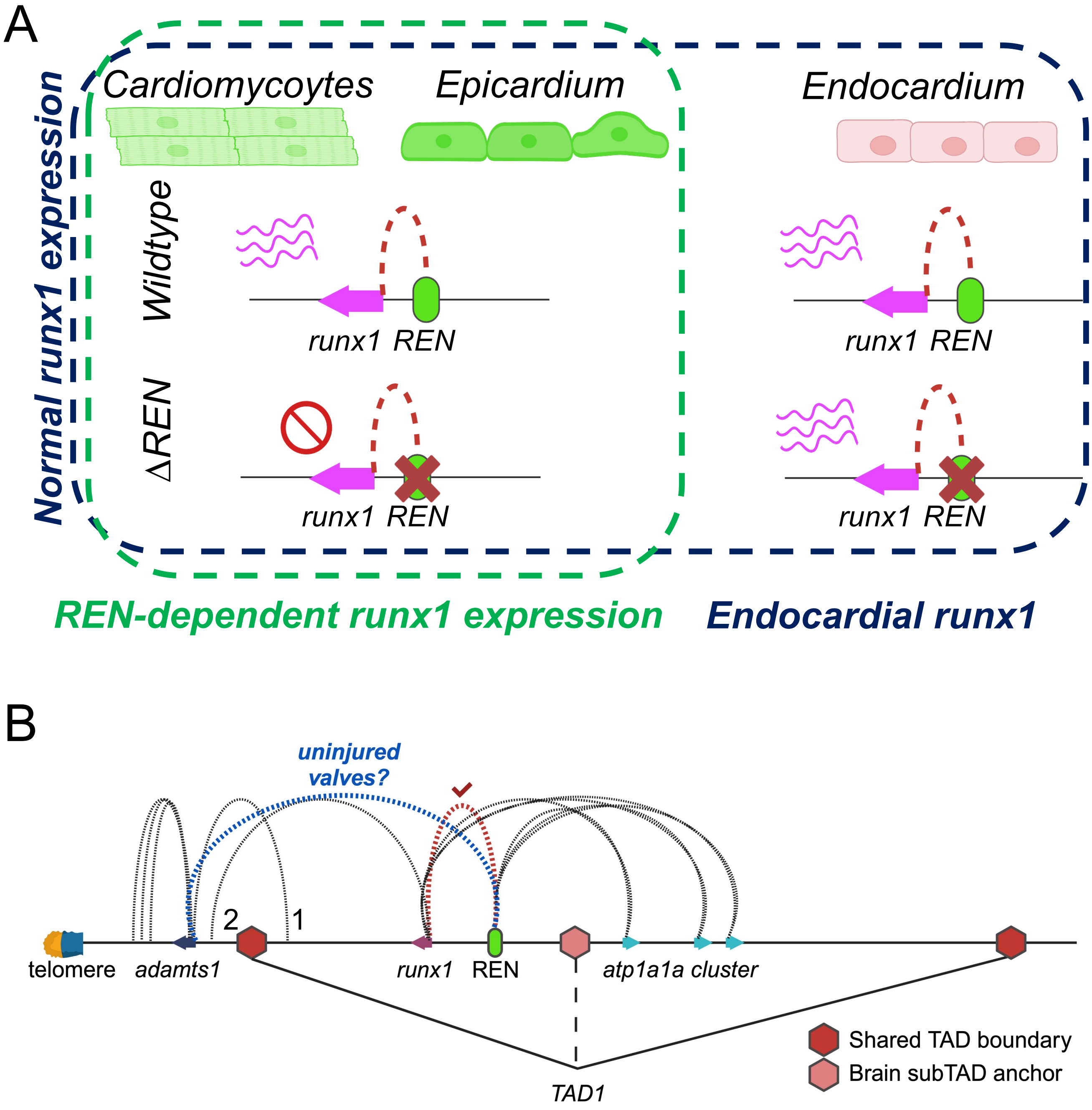
A
Normal runx1 expression
Cardiomycoytes
Epicardium
Endocardium
Wildtype
runx1 REN
runx1 REN
ΔREN
runx1 REN
runx1 REN
REN-dependent runx1 expression
Endocardial runx1
B
uninjured
valves?
2
1
telomere
adamts1
runx1
REN
atp1a1a cluster
TAD1
Shared TAD boundary
Brain subTAD anchor
